# Low-Cost Robotic Manipulation of Live Microtissues for Cancer Drug Testing

**DOI:** 10.1101/2024.03.21.586169

**Authors:** Ivan Stepanov, Noah R. Gottshall, Alireza Ahmadianyazdi, Daksh Sinha, Ethan J. Lockhart, Tran N.H. Nguyen, Sarmad Hassan, Lisa F. Horowitz, Raymond S. Yeung, Taranjit S. Gujral, Albert Folch

**Author notes:** These authors contributed equally to this work (Ivan Stepanov, Noah R. Gottshall).

## Abstract

The scarcity of human biopsies available for drug testing is a paramount challenge for developing new therapeutics, disease models, and personalized treatments. Microtechnologies that combine the microscale manipulation of tissues and fluids offer the exciting possibility of miniaturizing both disease models and drug testing workflows on scarce human biopsies. Unfortunately, these technologies presently require microfluidic devices or robotic dispensers that are not widely accessible. We have rapidly-prototyped an inexpensive platform based on an off-the-shelf robot that can microfluidically manipulate live microtissues into/out of culture plates without using complicated accessories such as microscopes or pneumatic controllers. The robot integrates complex functions with a simple, cost-effective and compact construction, allowing placement inside a tissue culture hood for sterile workflows. We demonstrated a proof-of-concept cancer drug evaluation workflow of potential clinical utility using patient tumor biopsies with multiple drugs on 384-well plates. Our user-friendly, low-cost platform promises to make drug testing of microtissues broadly accessible to pharmaceutical, clinical, and biological laboratories.

**Teaser:** A low-cost robot for handling microtissues and catalyzing their use in cancer drug evaluation and personalized oncology.

## Introduction

The process of drug development is vastly inefficient.^1,2^ The final stage of drug screening – a “drug evaluation” where the top drug candidates are tested for efficacy and safety to decide which candidate is pushed onto a clinical trial – is especially critical. The traditional disease models based on tissue cultures and animals are extremely poor predictors of human disease outcomes, as shown by decades of clinical trials.^3–7^ In oncology, where the first line of treatment is often surgery, an obvious solution to the poor predictivity of animal testing is to directly test drugs *ex vivo* on human tissue fragments or constructs. However, the typical size of a tumor biopsy extracted during surgery is only ∼1 cm^3^,^8^ which presents challenges for testing multiple drugs. Furthermore, the rise of combination therapies^9^ and dynamic therapies^10,11^ are exponentially increasing the complexity and cost of testing, and the improvement in early detection techniques^12^ keeps reducing the biopsy sizes obtained at the time of intervention.

As a result, there has been a fast-rising interest in miniaturizing the drug development process via the use of submillimeter-sized 3D live tissues (“microtissues”). These microtissues enable inexpensive, more efficient tests of high clinical biomimicry that address microscale phenomena (such as diffusive mass transport and the role of tissue heterogeneity on therapeutic efficacy^13^), while maximizing the use of scarce materials (*i*.*e*., live biopsies)^14,15,24–26,16–2^> and minimizing animal testing. The microtissues are produced either by bottom-up or top-down approaches. Bottom-up approaches include constructs such as organs-on-chips or organoids built by bioprinting,^27,28^ microengineering,^29–31^ or aggregation from single cells,^32–35^ often using cell growth to amplify the tissue. Conversely, the top-down approach consists of fragmenting biopsies by microdissection (without any growth), encompassing microdissected tissues such as organospheres, spheroids, and tumoroids.^15,16,25,26,17–24^ Microtissue-based drug tests are used not only for drug screening^16,17,35–37^ but, increasingly, also for disease modeling in cancer^15,38^ and immunology,^39^ for regenerative medicine,^40^ and for personalized medicine.^26,37,41^

Because of the microtissues’ small size, their high-throughput manipulation and culture has spurred the development of high-precision microfluidic and robotic platforms. Microfluidic tools such as hanging drops,^42–44^ droplet microfluidics,^26,2^> and/or microfluidic perfusion^11,15–17,21,34,45–48^ ensure the fluidic compartmentalization and microenvironmental control of the microtissues, however they entail complex fluid control systems that are not plug-and-play. Furthermore, microfluidic fabrication and operation require highly specialized human expertise, which can hamper their wider adoption in research settings and the translation of microtissues to the clinic. Various robotic platforms address the lack of user-friendliness of microfluidic platforms by automating the handling (*i*.*e*., pipetting and transferring) of microtissues and fluids. However, present robotic manipulators often target high-resolution manipulation of single cells requiring a microscope, a pneumatic controller, or both,^49^ and existing commercial systems typically integrate filtered air for sterility, resulting in bulky, expensive robots (see **Table S1**).

To broaden access to microtissue research, we have rapidly-prototyped a user-friendly, cost-effective and compact robotic platform that enables the automated manipulation of microtissues in a standard tissue culture hood (**Fig. 1** and **Fig. S1**). Compared to existing commercial robotic dispenser systems, our system features three key advantages: i) Instead of a microscope, a high-resolution USB camera (**Fig. 1**A) enables low-cost sorting and transfer of live microtissues. Aided by computer vision, the robot sequentially picks and sorts microtissues from a random distribution of microtissues in a culture dish into a multi-well plate (or any user-programmed array) (**Fig. 1**B). ii) Our platform builds on a compact off-the-shelf robot, a design that is compatible with standard sterile workflows (**Fig. S2**). Robotic placement of the microtissues into a multi-well plate in the hood is followed by culture in a standard incubator. iii) Fluid motion to manipulate tissues does not require a separate pneumatic controller or syringe pump; instead, the robot integrates a custom rotary pump powered by the motor in the end-effector (“head”) of the robotic arm (**Fig. 1**B inset & Rotary Pump in Suppl. Mater.). This pump generates the microfluidic flow needed to “lift” and dispense microtissues via a standard glass capillary. Thus, a single Python interface controls the pump via the translational motors, as well as the rest of the actions and user interface for the system. Hence, this integrated solution simplifies and substantially lowers the cost of programmable fluid manipulation.

**Figure 1.**
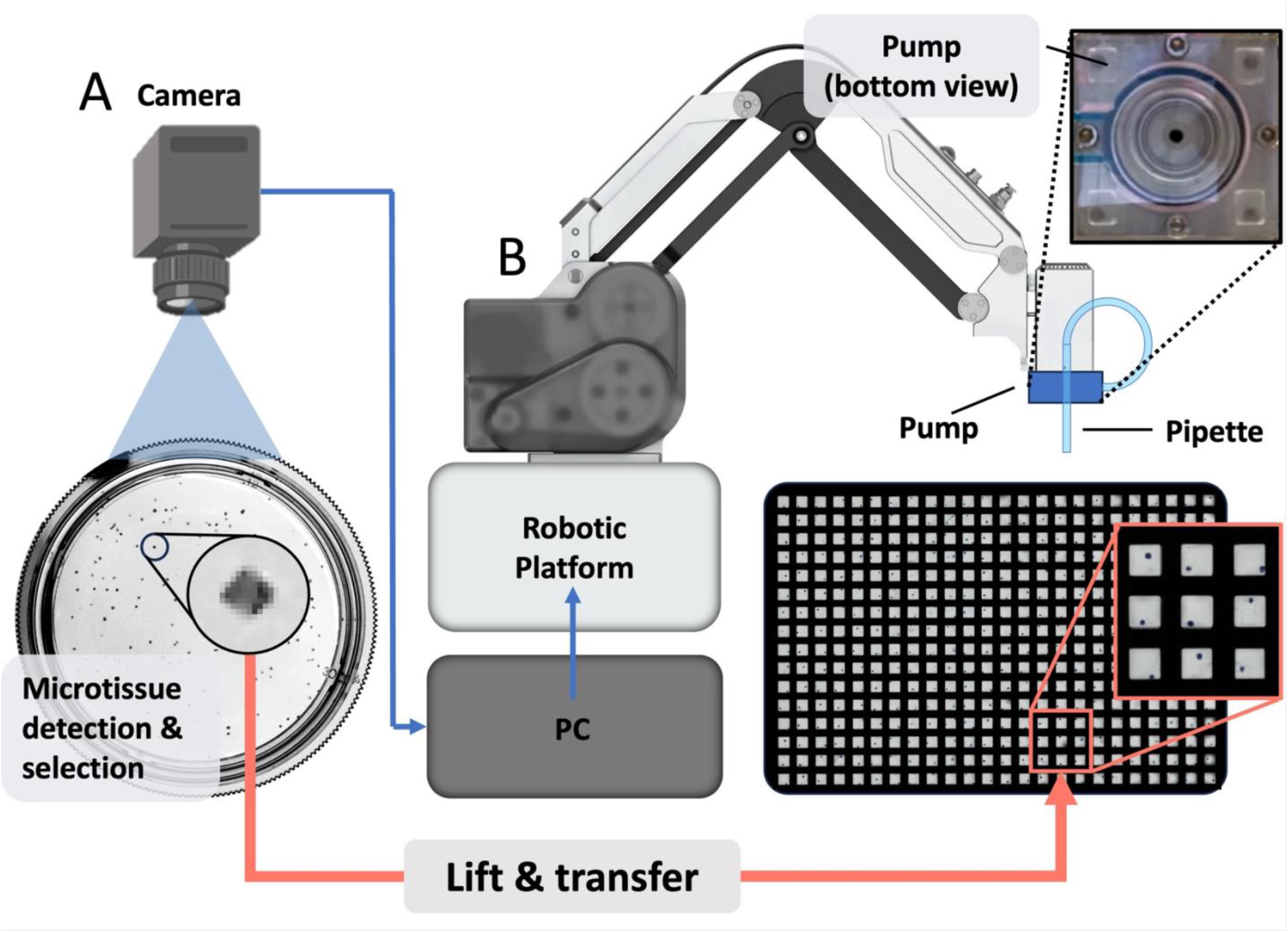
Robotic platform for automated ‘lift’-and-transfer of microtissues from a culture dish to a multi-well plate. **(A)** Schematic of the setup depicting the culture dish, a random distribution of microtissues (cuboids), and the USB camera above. Inset: magnified view of a 400 μm-wide cuboid illustrating that the resolution of the USB camera is sufficient to detect cuboids. **(B)** Conceptual rendering of the robot’s operation depicting the image that guides the robot’s movements, the robot assembled with the pump, and the capillary at its head above a 384-well plate (inset: 3 × 3 array of wells containing 400 μm-wide microtissues). The microtissue distribution is analyzed by a PC using images taken by the camera. The software selects a suitable microtissue to be transferred to the target well plate by the robotic platform. Activating the pump, the robot “lifts” the microtissue into the capillary and transfers it to the target well. Top inset: Bottom view of the rotary pump.

To demonstrate the utility of our platform, we present robotic manipulations that enable facile drug testing of monodisperse microdissected tissues (both mouse and human) in 348-well plates. These “cuboids” result from three orthogonal cuts with a tissue chopper (see Methods), featuring a relatively uniform distribution of sizes and cuboidal shapes at day 0,^16,4^> and retain the tumor microenvironment (TME).^50^ While shape uniformity is convenient, and the TME can play a critical role in drug efficacy, other types and shapes of microtissues such as minced tissues or organoids can also be used, and our freely-available GUI permits size selection. Human tumor cuboids usually retain their cuboidal shape for a few days in culture, whereas mouse tumor cuboids rapidly evolve to a spheroid shape in the same period, hence here we refer to the latter as “spheroidal cuboids” to avoid confusion, while the term “microdissected tissue” or “microtissue” applies to both. We show that the technology applies equally well for manipulating the tested microtissue shapes, sizes, and species. We demonstrate robotic protocols to pick and place single cuboids and sort hundreds of cuboids according to their size in separate wells in less than one hour. Finally, we perform drug evaluations of cuboids with multiple drugs on 384-well plates, including the evaluation of clinically-relevant drugs on a patient biopsy.

## Results

### Microtissue pick-and-place using a microfluidic ‘lift’-and-transfer process

Operation of the robotic platform is largely automated through a graphic user interface (GUI, see Methods). Through the GUI, the user calibrates the robot’s coordinate system (see Computer Vision in Suppl. Mater. and **Fig. S3**A&B), programs the destination positions for the microtissues (such as a multi-well plate), and specifies the range of microtissue sizes to pick from (see Computer Vision in Suppl. Mater. and **Fig. S3**C).

We evaluated the parameters that affect microtissue picking by our robotic platform. Using high-resolution video, we determined the accuracy and precision of the localization of microtissues. We measured the precision, or the repeatability of localization, as 26 μm ± 3 μm, which is well below the nominal 50 μm advertised by the robot manufacturer. We measured the accuracy, or the distance between the center of the capillary (after a given move) and the intended target (the center of the microtissue), as 129 μm ± 23 μm (**Fig. 2**A & **Movie S1**). Note that accuracy depends on the procedure for calibrating the position of the glass capillary, which is operator-dependent. While it could be improved, the accuracy of less than half the size of the microtissue size (250 or 400 μm^3^) proved to be sufficient.

**Figure 2.**
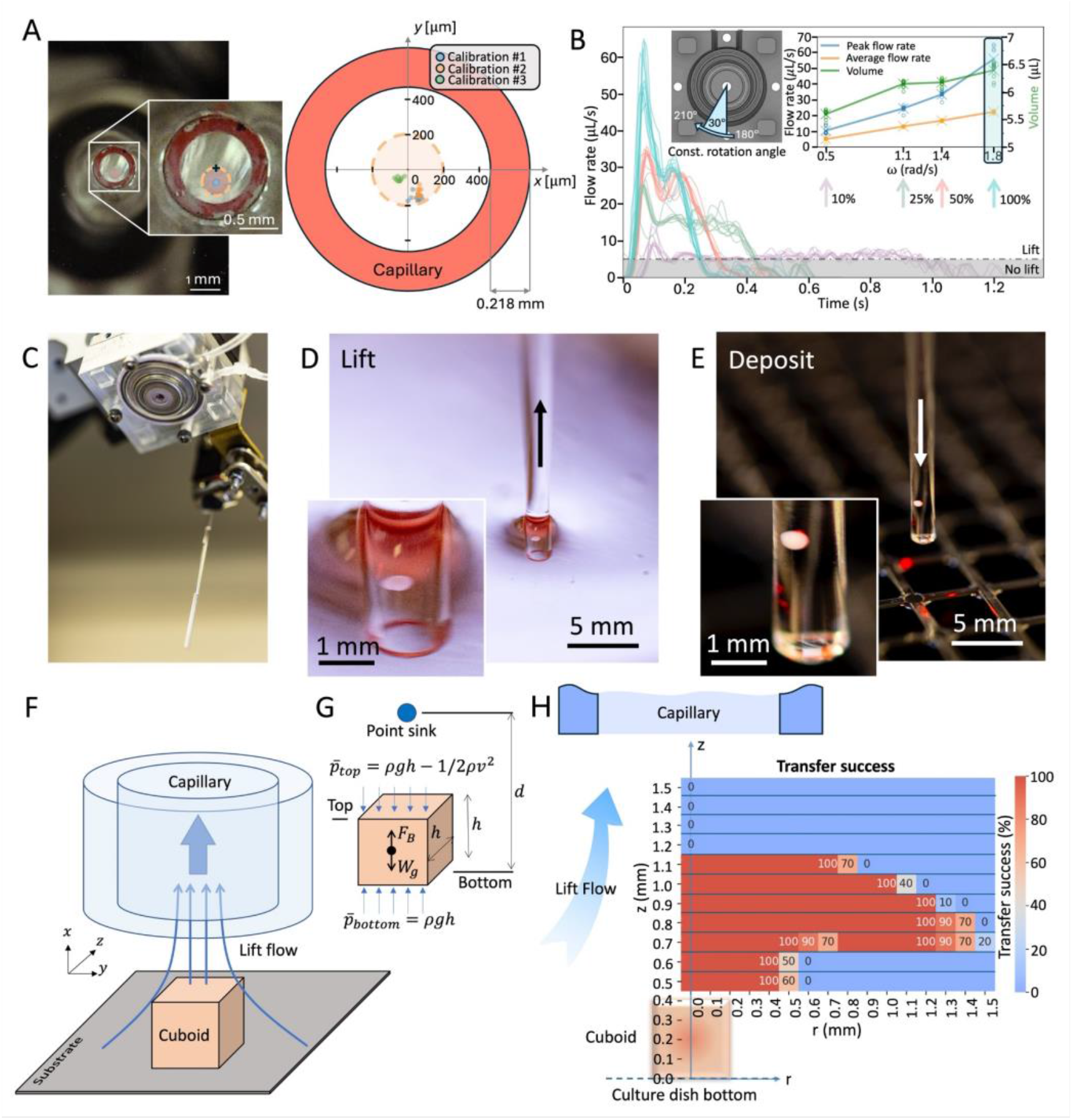
Microfluidic ‘lifting’ of microtissues. **(A)** Accuracy and precision of pipette localization to the tissue. (Left) Images of mouse cuboid (orange-dashed circle, blue centroid) taken from underneath the dish after approaching the capillary (red-painted rim). (Right) Graph depicting 30 attempts (10 attempts/calibration) to target the cuboid’s center (at the graph origin) with the capillary (shown as red torus for scale). **(B)** Plot of flow rate produced by pumping 30° rotations at four different angular speeds ω (100%, 50%, 25% and 10% of maximum ω, each plot consisting of ten repeats, colored as indicated by the vertical arrows). The black-dashed line indicates the lowest flow rate (∼5.1 μL/s) for which there is lift. (Left inset) Pump schematic displaying 30° rotation from 180° to 210°. (Right inset) Plot depicting maximum/average flow rate and volume pumped for the four ω values above. The cyan-shaded area indicates our operating regime. **(C)** Photograph of the pump and the capillary on approach to a culture dish. **(D)** Photograph of microtissue being lifted into the capillary. **(E)** Photograph of microtissue being deposited into a well. **(F)** 3D-schematic setup depicting the cuboid and the lift flow entering the capillary above it. **(G)** Schematic approximation of the capillary as a point sink, along with the main model parameters. **(H)** Graph depicting cuboid picking success (for a fixed PY8119 mouse breast cancer cuboidal microtissue) as a function of radial distance *r* from the cuboid’s center and the height *z* of the capillary above the culture dish bottom, with 2D schematics of cuboid (below) and capillary (above) to scale for reference.

To pick up live microtissues in culture medium from the culture dish surface, we used a microfluidic “lift” process (**Fig. 2**B-H) powered by a custom-made rotary pump. We rapid-prototyped the pump by CNC-milling PMMA plastic (see Methods) and installed it directly below the robot’s head. The head’s stepper motor aligns with and powers the rotary pump (**Fig. S1**B, **Movie S2**, and Rotary Pump in Suppl. Mater.). The rotary pump can thus be easily programmed via Python. We used a rollerless eccentric rotary pump design to avoid pulsatile flow delivery. The pump, which only pumps air, can rotate 360 degrees (or a fraction thereof) either backwards to generate suction or forward to generate positive pressure. The pump is connected to a glass capillary (VWR International, nominal I.D. 0.94 mm and O.D. 1.37 mm). A backward flow *Q* generates microfluidic “lift” that picks a microtissue off a surface (see Methods & **Movie S3**). The fluid at rest keeps the microtissue inside the capillary during the translation of the arm (neglecting the sedimentation speed of ∼500 μm/s). Forward flow dispenses the microtissue at the target location. Precise measurements of *Q*(*t*) as the pump rotates a fixed amount (30°) (**Fig. 2**B) were obtained with a flow sensor (Sensirion SLF3S-1300F) to determine *i*) the minimum *Q* that is needed to lift a microtissue off a surface (*Q*_*min*_ = 5.1 ± 0.3 μL/s, dashed horizontal black line in **Fig. 2**B inset), *ii*) the maximum *Q* that we can achieve (56 ± 6.1 μL/s using the highest rotational speed of the motor), and *iii*) the total volume entering the capillary by integration of the *Q*(*t*) curves (∼6.4 ± 0.1 μL at 56 μL/s). A large *Q* has the benefit of speeding up the process. **Fig. 2**C-E shows a sequence of three photographs depicting the pump and capillary approach to the culture dish, the capillary suctioning a microtissue (see **Movie S3**), and the moment when the microtissue is deposited into a well of a 384-well plate.

A theoretical fluid mechanics analysis illustrates why lifting a microtissue (in the diagram of **Fig. 2**F&G, a cuboid) does not require high-precision instrumentation (*e*.*g*., a microscope or, in its default, a proximity sensor) to position the capillary in *z* near the surface. For decades, biomedical engineers have automated the manipulation of droplets,^51^ cells,^52,53^ colonies,^54^ embryos,^55^ and worms.^56^ Electrostatic,^57^ magnetic,^58^ acoustic,^59^ and pneumatic soft^60^ end-effectors have been tethered to a robotic arm to manipulate small objects. For instance, Drury and Dembo^61^ examined the hydrodynamics of human neutrophils during micropipette aspiration, and Kumagai and Fuchiwaki^62^ developed a capillary dispenser that could pick-and-place objects. However, to the best of our knowledge, the physical process of how microscale flow can “lift” various small objects off a cell culture surface has not been described and validated quantitatively. In our setup, the capillary mounted at the head of the robot provides the “lift” flow (**Fig. 2**F) that we can approximate as a point sink (**Fig. 2**G). For a cuboid of volume *V*_*c*_ and density *ρ*_*c*_, its weight is *W* = *ρ*_*c*_ *V*_*c*_*g* and the buoyant force it experiences is *F*_*B*_ = *ρV*_*c*_*g* where *ρ* is the density of the fluid (water). Hence the following inequality must be satisfied:

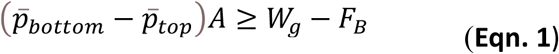

where 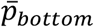 and 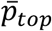 are the average fluid pressures at the bottom and top surfaces of the cuboid, respectively, and *A* ∼ (400 μm)^2^ is the area of each of the faces of the cuboid. Applying Bernouilli’s equation, 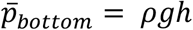 and 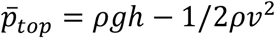, where *ν* is the average flow velocity on top of the cuboid, **Eqn. 1** can be re-written as:

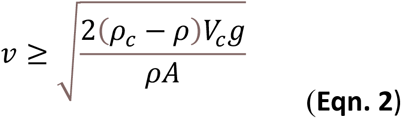

Since *ν* has components *u*_*x*_, *u*_*y*_, and *u*_*z*_, where *u*_*y*_ and *u*_*z*_ ≪ *u*_*x*_ because the capillary is centered with the cuboid, then *ν* ≈ *u*_*x*_, where *u*_*x*_ can be obtained from ideal flow equations through the method of images:^63^

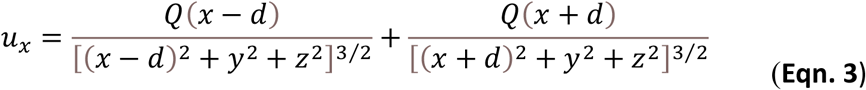

In **Eqn. 3**, *Q* is the instantaneous suction flow rate generated by the pump (as measured by our flow sensor), *d* is the distance between the cuboid’s bottom surface and the point sink, and *x, y*, and *z* refer to the distances from the coordinate system located at the cuboid. Thus, we can find the flow velocity on top of the cuboid by substituting *x* = *h* into **Eqn. 3**. Assuming *h* ≪ *d*, we obtain *ν* = 2*Qh*/*d*^3^. Substituting into **Eqn. 2**, we obtain a condition for cuboid lifting to occur:

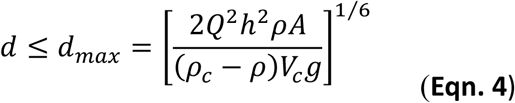

where *h* ≈ 400 μm. A similar analysis follows from assuming a spheroidal shape. Note that *d*_*max*_ is not very sensitive to changes in any of the parameters. Since *A* ≈ *h*^2^ and *V*_*c*_ ≈ *h*^3^, then *d*_*max*_ ∼ *Q*^1/3^*h*^1/6^. Substituting our typical values (*Q* = 56 μL/s, *ρ*_*c*_ = 1.079 g/cm^3^ for mouse tumors^64^), we predict that the maximum distance between the capillary and the bottom of a cuboid that allows for lifting the cuboid is *d*_*max*_ ≈ 1.2 mm for a 400 μm cuboid and *d*_*max*_ ≈ 1.1 mm for a 250 μm cuboid.

Measurements of picking success as a function of *z* for 400 μm cuboids (**Fig. 2**H & **Fig. S4**) validate this analysis. For these experiments we used fixed PY8119 mouse breast cancer spheroidal cuboids. In agreement with our simplified point-sink model, we find that the capillary needs to be equal or closer than ∼1.1 mm above the surface to lift a 400 μm mouse tumor cuboid. We note that the model assumes that *Q* is applied as a step function, but the peak shape of *Q* could help lift the cuboid as soon as it surpasses *Q*_*min*_ ≈ 5.1 μL/s (**Fig. 2**B), making the effective value of *d*_*max*_ larger. In sum, the microfluidic lift effect occurs for a wide substrate-to-capillary distance range of 0.4-1.1 mm (0-700 μm above a 400 μm cuboid), which in practice makes a microscope or a proximity sensor unnecessary.

### Error correction

The robotic platform incorporates software-based checks and diagnostics that prevent and correct errors in cuboid manipulation. Two main types of error can occur: an empty well or a well filled with two microtissues. To prevent the possibility of lifting two or more adjacent microtissues and determine the minimum empty distance surrounding a target cuboid, we measured the sensitivity of the capillary for picking a nearby microtissue. We first measured the success of picking a 400 μm cuboid as a function of *r*, the lateral distance between the center of mass of the cuboid and the center of the capillary, for many values of *z* (**Fig. 2**H). The transfer success plot displayed an interesting “mushroom” profile: right above the cuboid (within 0.4 mm, or its own height), the capillary only lifted the cuboid when its center was within the margins of the cuboid. However, at heights larger than 0.6 mm above the cuboid, the lift radius L_R_ from the center of the cuboid increased suddenly from L_R_ = 0.4 mm to L_R_ = 1.2 mm (**Fig. 2**H). At the usual *z* = 0.8 mm where we placed the (end of the) capillary, the success decayed rapidly to 0% for *r* > 1.5 mm. This mushroom profile has an immediate consequence on the ability to discriminate between two adjacent cuboids: the user can either choose to hover at low capillary-to-substrate *z* heights (risking crashing of the capillary) for high selectivity or hover at higher *z* (thus losing the ability to discriminate between two adjacent cuboids) for higher safety. We decided on higher safety because not all culture dishes are equally planar, and it is straightforward to use image recognition methods to select for properly-distanced microtissues. Note that both the lateral and height spread of the red “mushroom” are very similar for the cuboid and spheroid shapes (**Fig. S4**), indicating that the lift mechanism does not depend strongly on microtissue shape. Hence, for safety, we operated at *z* = 0.8 mm and picked only microtissues that are at least 2 mm away from another microtissue (see Transfer Success in Suppl. Mater. & **Fig. S4**).

To prevent picking up two microtissues at once, the software scans the culture dish and measures the distance between each microtissue and its nearest neighbor, allowing the user to filter out microtissues that are critically close to another microtissue (see Computer Vision in Suppl. Mater., and **Fig. S3**D). However, we observed that, in the rare cases when two small microtissues are adhered to each other, they may be interpreted as a singular, normal-size microtissue that is brought into the target well together but separates into two after ejection. Moreover, if a microtissue adheres to the inner wall of the glass capillary, its failure to eject is counted as an empty well; when the next microtissue is lifted, the microtissue buildup inside the capillary can dislodge the previously-stuck microtissue, resulting in a well filled with two microtissues. In human cuboid cultures, which tended to aggregate more strongly than mouse spheroidal cuboids (likely in part due to our use of serum-free, defined medium), more than half (∼56 ± 6.1%) of the wells that were filled with two cuboids were also preceded by empty wells. In mouse cuboid cultures, on the other hand, wells filled with two cuboids were never preceded by empty wells, which confirms that errors can depend on tissue type or shape.

We also addressed empty wells. Most empty wells tended to be caused by the robot’s physical inability to pick up certain microtissues, *e*.*g*., because the microtissues stick to the bottom of the culture dish, or conversely, they were not sufficiently attached during the capillary’s approach and tended to float. Since empty wells can be corrected with a second attempt, to rectify empty well errors we used a corrective algorithm consisting of comparing images taken before and after the picking attempt (see Computer Vision in Suppl. Mater.). If the microtissue is still in the culture dish in the same position where it was before picking, the robot returns the (microtissue-free) liquid back into the culture dish and tries again. Since the liquid ejection modifies the original distribution of microtissues at that particular location, the robot resets and attempts to pick anew. Note that our platform could not pick occasional, rare microtissues that float (*e*.*g*., those that have a high fat tissue content or have attached air bubbles) because they can easily change their position due to liquid movement after the pickup attempt. Likewise, occasional bubbles present in the culture dish can cause an empty well, but preventive measures in the software mainly take care of them (see Computer Vision in Suppl. Mater. and **Fig. S3**D).

To obtain statistics on the success rates of microtissue transfer by the robotic platform, we filled six 384-well plates with microtissues from different batches: two well plates were filled with Py8119 mouse tumor cuboids, and four well plates were filled with two colorectal cancer (CRC) liver metastasis human tumor cuboids. We defined success as a well filled with a single microtissue. For mouse cuboids, we observed that 98.4 ± 0.2% of the transfer attempts were successful, 0.5 ± 0.0% were empty wells, and 1 ± 0.2% were two cuboids instead of one. Human CRC cuboids were somewhat more difficult to lift and dispense (appearing “stickier”), resulting in an average compound success rate of ∼92 ± 1.6%, 4.3 ± 0.9% with empty wells, and 3.7 ± 0.7% with doubles.

### Selecting microtissue size and transferring multiple microtissues at once

Microtissue size can be a confounder in studies such as cytokine secretion (the number of cells in the microtissue affects the secretion readout) and hypoxia-induced cell state (the microtissue size determines the oxygen diffusion path length, and low oxygen can lead to growth arrest or even death). The image resolution and size determination algorithm are accurate to within ± 3.5 μm (see Computer Vision in Suppl. Mater.). To reduce variability in size, we implemented a size-sorting software feature that allowed us to pick microtissues with different narrow microtissue size ranges, followed by delivery to pre-programmed areas of a multi-well plate (**Fig. 3**A). These experiments demonstrated size ranges between 50 and 130 μm with fixed mouse tumor cuboids. The setup is not restricted to 400 μm-wide microtissues; a simple change of the size threshold in the software can also enable other applications which call for the manipulation of much smaller and larger microtissues.

**Figure 3.**
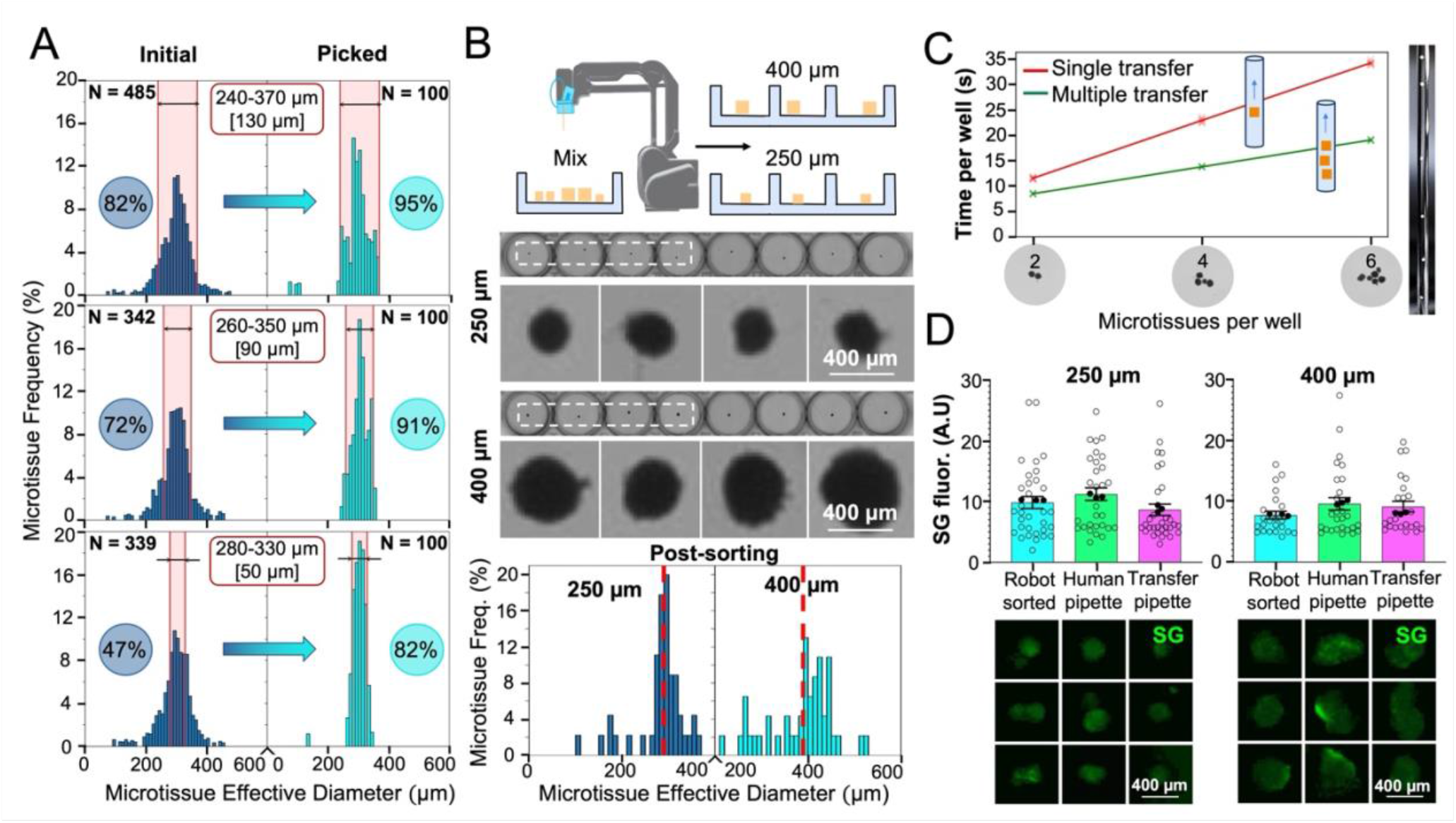
Robotic size-sorting of microtissues. **(A)** Three consecutive experiments for size selection of fixed Py8119 mouse breast cancer cuboids with a decreasing picking size range from top to bottom: 130, 90, and 50 μm, respectively. The few data points below 200 μm represent camera sensor noise and debris in the culture dish. **(B)** Robotic sorting of a mixture of 250 and 400 μm-diam. live Py8119 cuboids into a 96-well plate. (Top) Process schematic. (Center) Grayscale micrographs of two rows of the 96-well plates after sorting and filling with 250 and 400 μm-diam. cuboids. (Bottom) Post-sorting size histograms. **(C)** Graph of robotic transfer duration as a function of number N of cuboids per well (depicted in X axis with representative images), comparing the transfer of N cuboids one at a time (red line) or N cuboids at once (green line). (Right) Photo of 6 cuboids loaded into the capillary. **(D)** Absence of viability loss in live Py8119 cuboids due to robotic transfer. (Top) Graphs of SYTOX Green (SG) fluorescence (cell death) comparing three transfer methods (robotically transferred, manually transferred with a pipette, and manually transferred with a transfer pipette) for ∼250 and ∼400 μm cuboids. Average ± sem, each point represents a cuboid. 400 μm cuboids: n = 25-30/condition; 250 μm cuboids: n = 32-35/condition. Kruskal-Wallis test with Dunn’s multiple comparisons test yielded no significant statistical differences for both size groups. (Bottom) Close-up micrographs of three spheroidal cuboids representative of the mean SG fluorescence (black-filled points in the graph) for each condition.

We demonstrated accurate size-selective sorting from a random mixture of 250 and 400 μm-diam. live Py8119 mouse breast cancer cuboids (**Fig. 3**B). The post-sorting size distributions of the separated populations of 250 and 400 μm spheroidal-shaped cuboids revealed an average size of 288 and 391 μm, respectively (red dashed vertical bar in **Fig. 3**B graph). The robot can also pick more than one cuboid (up to six) at once to save time if loading a well with multiple cuboids is desired; as shown in **Fig. 3**C, multi-cuboid transfer was 1.35×, 1.66× and 1.8× faster for 2, 4, and 6 cuboids, respectively, than the added time of transferring each cuboid separately.

To ensure that robotic transfer does not interfere with tissue viability, we compared transfer of live Py8119 cuboids (250 and 400 μm) by robot (with ∼1 mm I.D. glass capillary tube) with two standard hand-operated pipette-based methods: a P200 pipette (with tip cut to ∼1 mm I.D., used for precise individual cuboid transfer) and a transfer pipette (∼1 mm I.D., a gentle method also used for all bulk cuboid preparation and transfer processes). We measured cell death with the fluorescent-green nuclear cell death indicator SYTOX Green (SG) 1 hour after transfer. We found no significant statistical differences between the three methods, indicating that robotic transfer causes no more significant loss of viability to the microtissues than transfer with the standard pipette-based methods (**Fig. 3**D).

### Drug tests with the robotic platform

Mouse cuboids, with their relative homogeneity and availability, allowed us to easily test the use of the robotic platform for multiplexed drug evaluations in multi-well plates (**Fig. 4**). We used the robot to load U87 glioma mouse xenograft tumor spheroidal cuboids (250 and 400 μm-diam.) from two different 6 cm culture dishes into separate 96-well plates (∼15 min / 96-well plate; see **Movie S4**). We measured cell death with SG fluorescence at the end of the 3-day drug treatment. The drug set included cisplatin (CP; DNA/RNA synthesis inhibitor), bortezomib (Bort; proteasome inhibitor), mocetinostat (MOC; HDAC inhibitor), parthenolide (Parth; NF-κβ inhibitor), YM155 (YM; E3 ligase/surviving inhibitor) and tanespimycin (AAG; heat shock protein inhibitor) at two concentrations for each drug, as well as dimethyl sulfoxide (DMSO; vehicle control for all but CP) and medium alone (control). The fluorescence readouts for both 250 and 400 μm cuboid plates revealed similar strong drug responses for AAG, MOC, Bort, and CP, as well as for the cell death control, staurosporine (STS). There were also differences. For Parth, both sizes appeared to respond, but only reached significance for 400 μm. For YM, which generated highly variable responses in the cuboids, only the highest concentration with 250 μm cuboids reached statistical significance. This experiment demonstrated the suitability of the robotic platform for drug treatments using both 250 μm and 400 μm cuboids.

**Figure 4.**
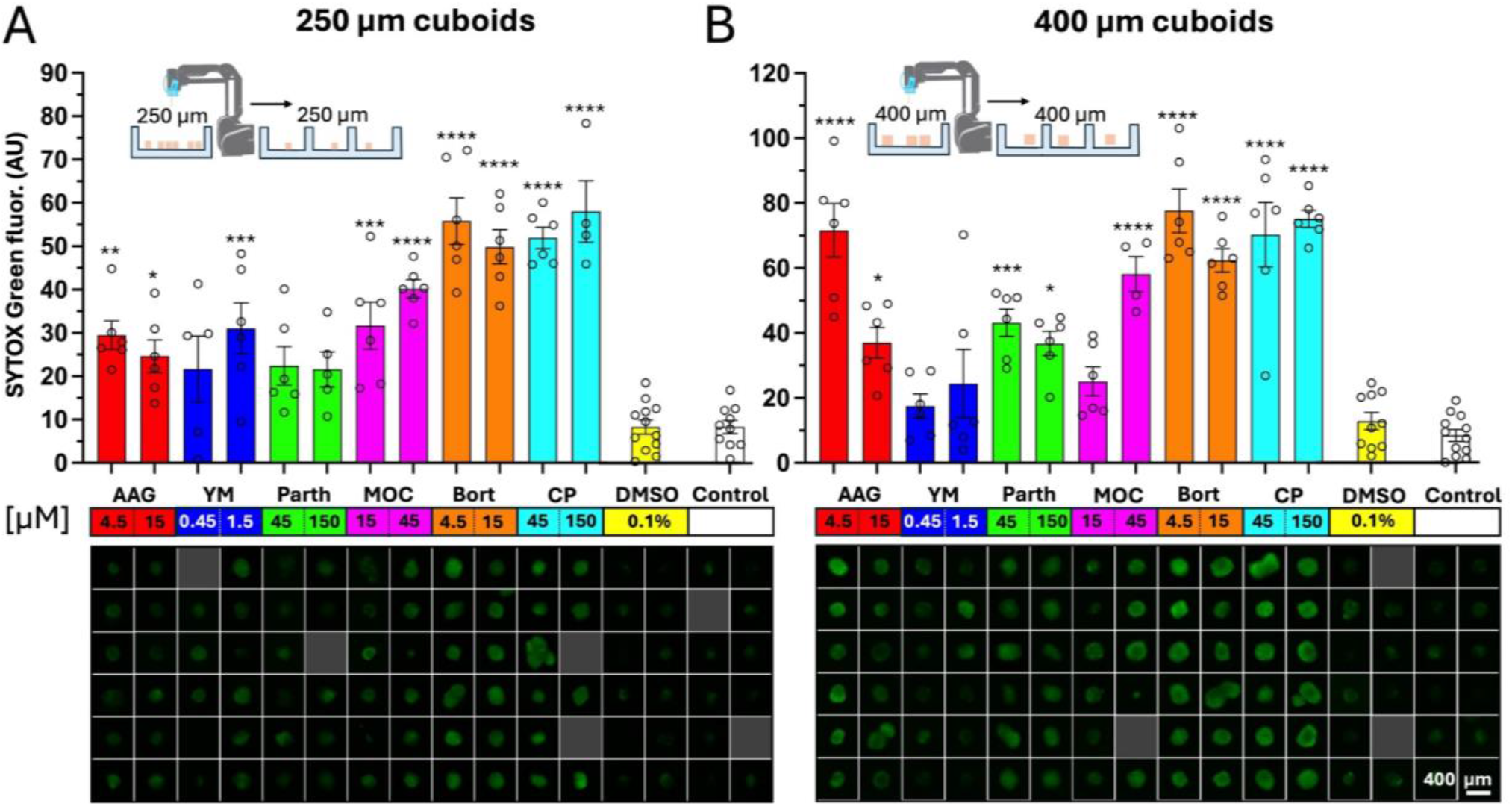
Drug testing of mouse tumor cuboids in 96-well plates. Drug treatments of 250 μm diam. (**A**) and 400 μm diam. (**B**) U87 mouse tumor cuboids. (Top) Bar graphs of SYTOX Green (SG, cell death) fluorescence readouts for each condition after 3 days of drug treatment. Separate batches of individually cut and segregated 250 and 400 μm cuboids were transferred into two different 96-well plates. (Bottom) Close-up micrographs of representative wells for each condition displaying green channel SG fluorescence. Each plate was treated with the following panel of drugs at two concentrations for each (drug concentration shown in μM): dimethyl sulfoxide (DMSO; vehicle control), cisplatin (CP; DNA/RNA synthesis inhibitor), bortezomib (Bort; proteasome inhibitor), mocetinostat (MOC; HDAC inhibitor), parthenolide (Parth; NF-κβ inhibitor), YM155 (YM; E3 ligase/surviving inhibitor) and tanespimycin (AAG; heat shock protein inhibitor). Average ± sem, each point represents a cuboid. n = 4-6 cuboids per condition. One-way ANOVA with Tukey *post-hoc* versus DMSO, except for CP (versus control). *p < 0.05, **p < 0.01 ***p < 0.001, ****p < 0.0001.

To evaluate the reproducibility of the cuboid drug treatment assay, we repeated the experiment on a different U87 mouse tumor, with 400 μm cuboids only on duplicate plates (**Fig. S5**). We treated the cuboids with the same drug panel at similar concentrations (one low and one high). The cell death SG fluorescence readouts were similar between the two 96-well duplicate plates. Combining the results of the duplicate plates to increase our sample size, we observed statistically-significant responses with AAG, Bort, CP, and STS, but not with YM or MOC, both of which showed increases but with high variability. These results were very similar to the results seen on the 400 μm cuboids from a different tumor (**Fig. 4**), supporting the reproducibility of the cuboid drug test.

With the robotic platform, scaling up the drug tests to 384-well plates was straightforward, assisted by halving the transfer volume to 5 μL. We performed a proof-of-concept two-drug dose response treatment on Py8119 mouse breast cancer cuboids in 384-well plates, in which we treated for 3 days with CP and STS at five different logarithmic concentrations (**Fig. 5**). The use of 384-well plates allowed for ∼32 cuboids per drug condition and demonstrated the robot’s ability to handle large sample sizes efficiently. In this experiment, the robot filled the plate in ∼65 min with a success rate of 98.4% (378 wells of 384 filled with one cuboid); 1% (4/384) of the wells were filled with two cuboids, and 0.6% (2/384) were empty. The SG fluorescence cell death readouts for each condition revealed cell death relative to control cuboids for the two drugs, with a trend of nearly linear increasing cell death at increasing STS concentrations.

**Figure 5.**
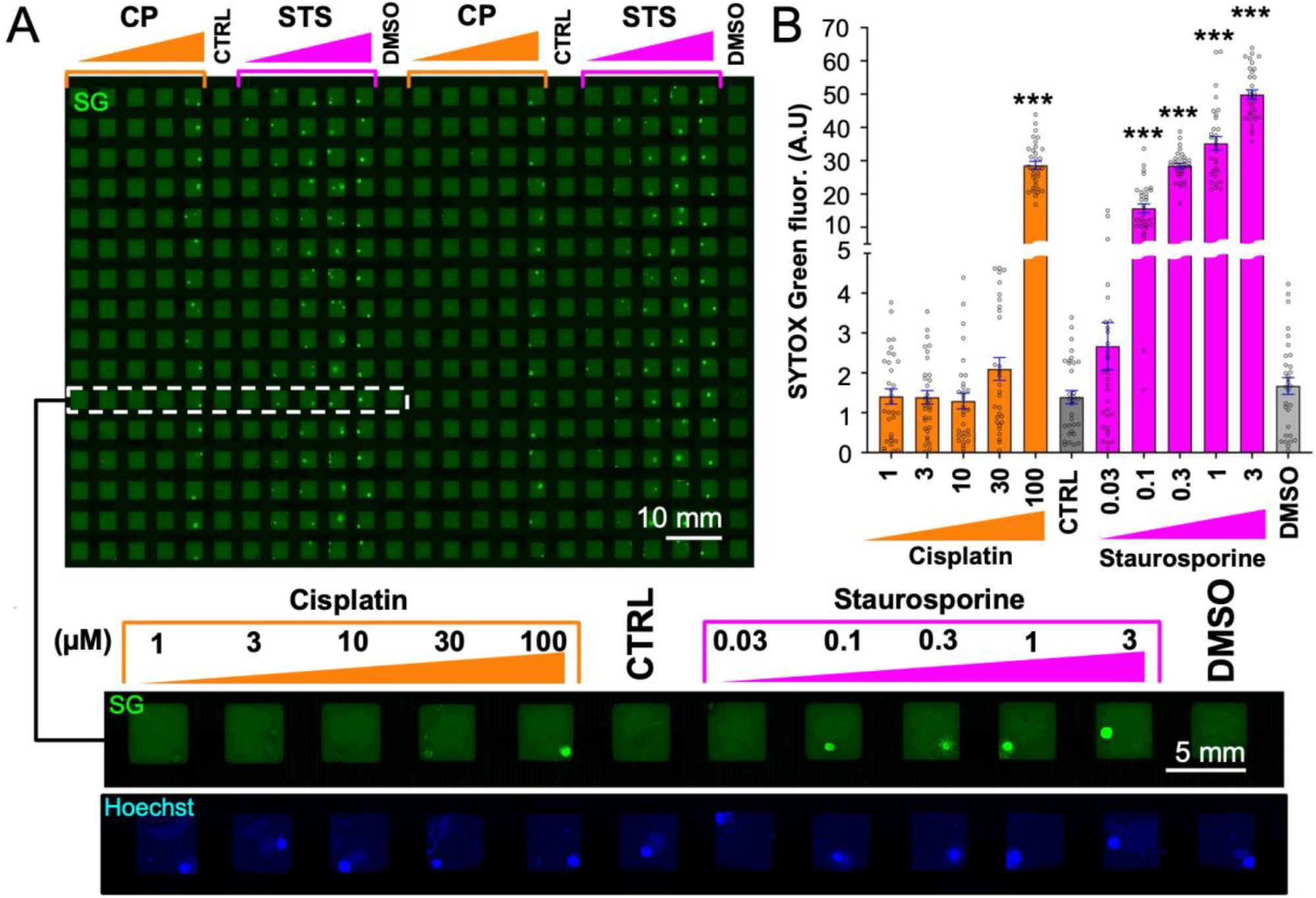
Drug testing of mouse tumor cuboids in a 384-well plate. **(A)** Py8119 mouse breast cancer spheroidal cuboids stained with SYTOX Green (SG) cell death stain in a 384-well plate after 3-day drug treatment. (Top) Green channel image displaying (visually) increasing fluorescence intensity with increased drug concentration. (Bottom) Close-up micrographs of blue and green channels for one complete treatment row of cisplatin (CP) and staurosporine (STS) at five different concentrations each (Hoechst nuclear counterstain (blue) used for cuboid identification). **(B)** SG fluorescence readout of 384-well plate for each condition. Average ± sem, each point represents a cuboid. One-way ANOVA with Tukey post-hoc test versus control (CTRL, no drug) for CP and DMSO (0.1%) for STS. n = 30-32 cuboids per condition. ***p < 0.001.

### A drug evaluation with human tumor cuboids

The simplicity of our robotic platform is ideal for performing direct drug evaluations on human tumor cuboids in a clinical context. Towards that end, we simulated a personalized oncology drug test with clinically relevant drugs using our robotic workflow and cuboids from a patient’s colorectal cancer (CRC) liver metastasis. The test took just over a week, yielding results rapidly enough, in principle, to influence treatment decisions. The patient was a 54-year-old male with recurrent CRC after previous surgery, infrared liver ablation, and chemotherapy with a combination of cisplatin, irinotecan, leucovorin and 5-fluorouracil (5-FU). We extracted three 6-mm core biopsies and prepared cuboids from six 400 μm-thick slices for each of the three cores (**Fig. 6**A). Using the robot, we filled the wells of three 384-well plates with individual cuboids, with one plate for each of the three cores, yielding an overall success rate of 91.1%. (Out of the 1,046 wells total for the three plates, the robot filled 953 wells with single cuboids, 41 wells with two cuboids, and 52 wells were empty.) The filled 384-plates allowed for 30-43 wells per condition. Before the drug test, we measured the baseline viability after overnight culture (**Fig. 6**B). The baseline viability between cuboids was much more variable than in mouse, as may be expected from a heterogeneous patient tumor; as apparent in the slices used to make cuboids (**Fig. 6**C), some of the darker areas appeared to correspond to acellular, stromal areas.^65^ Furthermore, baseline viability differed between cores, with the highest viability from core 1, followed by core 3 and then core 2.

**Figure 6.**
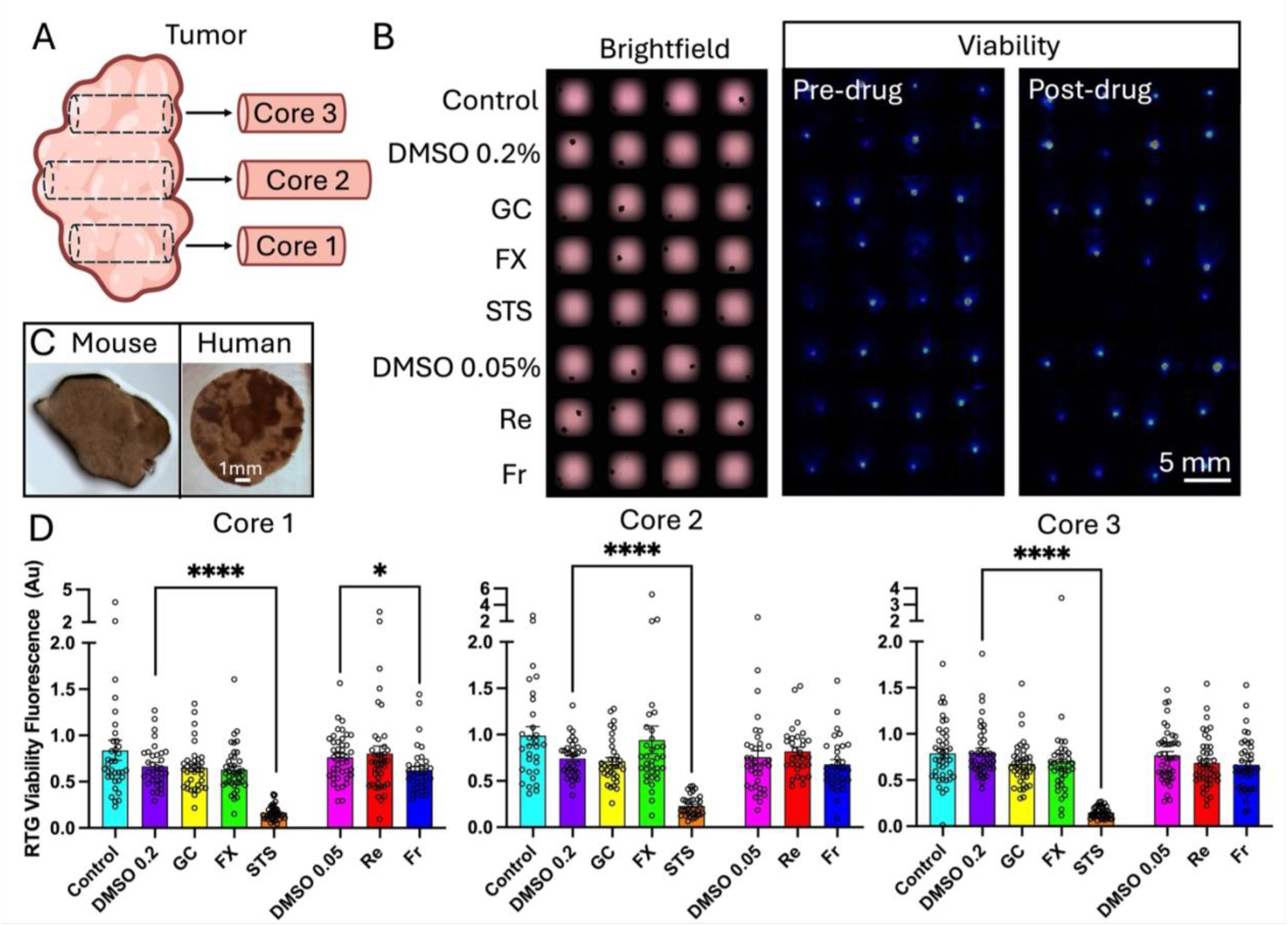
Drug testing on cuboids from a patient colorectal cancer liver metastasis with clinically relevant drugs. **(A)** Schematic of tissue preparation: three core biopsies were taken from the tumor and cuboids from each core were distributed onto separate 384-well plates. **(B)** Example region from the 384-well plate filled with cuboids from core 3. (Left) Bright-field micrograph displaying cuboid positions within wells, with each row corresponding to one condition. Conditions: DMSO (dimethyl sulfoxide 0.2 or 0.05 μM; vehicle control), a mixture of gemcitabine and cisplatin (GC; chemotherapy drugs); FOLFOX (FX; like the combination chemotherapy regimen for CRC with 5-fluorouracil and oxaliplatin); staurosporine (STS; broad kinase inhibitor used as a positive control); regorafenib and fruquintinib (Re and Fr, respectively; kinase inhibitors used for CRC). (Middle) Baseline viability of the same cuboids as luminescence measured with an IVIS machine. (Right) Post-drug addition viability measured on the 6^th^ day. **(C)** Examples of a mouse tumor slice (homogeneous) and human tumor slice (heterogeneous). **(D)** Graphs of the drug response ratios (day 6 / day 1) displayed for each core/plate. Ave ± sem, each point represents a cuboid. One-way ANOVA with Tukey post-hoc. **** p < 0.0001, * p < 0.05. n = 30-43 wells per condition.

To test drug response, we picked four clinical drug treatments which could be considered as the next therapy for this patient. FOLFOX (a combination of folinic acid, 5-FU and oxaliplatin), and gemcitabine/cisplatin (GC, a combination of gemcitabine 1 μM and cisplatin 5 μM) are standard cytotoxic chemotherapies for CRC liver metastasis; regorafenib (Re, 0.5 μM) and fruquintinib (Fr, 0.5 μM) are targeted kinase inhibitors approved for CRC.^66,67^ In addition, STS (1 μM) was used as a positive control. DMSO vehicle controls (0.2% for cytotoxic drugs, 0.05% for kinase inhibitors) as well as a medium alone served as negative controls. We measured viability by RTG at baseline before drugs, then after 4 days for the faster acting cytotoxic drugs, and after 6 days of drug exposure, as some drugs may have a slower action (**Fig. 6**B). RTG, unlike the single-time-point fluorescent SG assay, allows multiple readings of cell death and is not affected by the high autofluorescence often found in human tissues. Cuboids from all three cores showed the expected strong response to STS (decrease of 75 ± 7%, n = 3 cores), and showed no statistically-significant response to FOLFOX or GC (**Fig. 6**D). The lack of a statistically-significant response to FOLFOX and GC may be in part explained by resistance from the tumor, because the tumor had progressed before resection after the patient was treated with some of the compounds (5-FU in FOLFOX and cisplatin in GC). Of note, core 1 showed a response to Fr (decrease of 18.4%, p < 0.05) while the other two cores only showed a trend that did not reach significance.

Whereas the similarity between the drug response graphs corresponding to each core speaks for the high reliability of our assay, the ability to discern small differences between cores supports that the assay can also detect differences due to heterogeneity between different regions within a patient’s TME. Furthermore, individual cuboids in separate wells also provided a measure of the spatial heterogeneity of the patient tumor within a smaller, sub-millimeter scale. The assay’s sensitivity and the number of cuboids required to detect a given effect are influenced by the tumor’s heterogeneity, here seen as differences between individual cuboids in separate wells. We used a *post-hoc* power analysis to suggest the minimum sample size needed to see a certain effect with 80% power and 0.05% statistical confidence, given the variance in the values seen in this real-world test. To detect an almost complete response (*e*.*g*., the observed 75% decrease with STS), we would need to test only 2 cuboids per condition. On the other hand, to detect a very weak effect (*e*.*g*., the statistically-significant 18% decrease with Fr in core 1), we would need to test at least 55 cuboids per condition. For an intermediate effect (*e*.*g*., a 50% decrease), we would have needed 5, 12, or 6 cuboids per condition, estimated from the variability seen for cores 1, 2, and 3, respectively.

In sum, the robotic cuboid platform allowed us to rapidly test for differences (and similarities) in the drug response both regionally (between cores) and locally (cuboid to cuboid), highlighting the heterogeneity of the TME. The non-destructive viability assay and the standard well-plate format facilitates deeper analysis of the cuboids and their TME, *e*.*g*., by cytokine secretion or –omics.^50^ Hence, we believe that robot-enabled cuboid experiments could help jump-start TME-dependent research into cancer and its treatment, as well as cancer drug development.

## Discussion

Our robotic platform aims to catalyze microtissue research and its applications by providing a low-cost approach to microtissue manipulation. The platform has advantages and limitations compared to existing robotic dispensers and microfluidic devices. Like other robotic approaches, the GUI does not require specialized well plates and is not limited to regular arrays for drug testing, as it could be programmed to load other types of microtissue-sensing devices.^68,69^ The microfluidic “lifting” mechanism is universally applicable to any microtissue, as long as the microtissue fits in the capillary and the pump’s flow rate is high enough to counterbalance the weight of the microtissue. Although this study was focused on microdissected tissues, we demonstrated that the computer vision and lifting mechanism work equally well for cuboidal and spheroidal microtissues (**Fig. 2**H and **Fig. S4**, respectively), strongly suggesting that the platform would work for other microtissue formats such as organoids or organospheres that only differ in density or size since *d*_*max*_ is insensitive to changes in these parameters (**Eqn. 4**), i.e. *d*_*max*_ ∼ (*ρh*)^1/6^. In principle, the pump’s physical and operational parameters can be modified to produce much larger flow rates and in larger capillaries than shown here if much larger microtissues needed to be lifted. Compared to pneumatic pipette dispensers, which contain expensive components, our compact robot can be built with low expertise and very cost-effective parts to fit in a standard tissue culture hood. Our proof-of-concept system lacks multi-pipetting and fluorescence imaging capabilities present in most robotic, microscope-based systems, but improvements in the camera setup (*e*.*g*., fluorescence imaging, now performed offline) could add further capabilities to our platform. Arguably, articulated robots such as ours have complex kinematics, so they are not very fast, but less complex and faster (but less cost-effective) gantry-type robots will likely achieve similar pick-and-place operations in less time and could incorporate multi-pipetting fluid dispensing, potentially opening the way for multiple 1,536-well plate tests. Compared to microfluidic devices that can trap microtissues in a multi-well format in parallel,^16,4^> our robotic platform only manipulates microtissues serially (thus slower) and cannot generate microenvironments (*e*.*g*., as needed for microvascular studies^35,40,44,47^). However, in our platform, speed is traded for reliability, user-friendliness, and automation; in contrast to microfluidic channels which have a fixed design and have the risk of errors due to clogging, our robotic platform can be flexibly programmed to transfer microtissues into different layouts and has been designed to detect and correct its errors. While the corrective and preventive measures in place do not completely prevent errors from happening, they significantly reduce the error rates. The small amount of failure rates remaining should not pose a problem for drug test analysis, as the failures can be easily discarded during imaging.

The robotic cuboid platform is of general applicability to drug testing, especially for TME-sensitive drugs such as immunotherapies. Hence, it could be used to develop advanced intact-TME cancer models^50^ and, with the help of machine-learning algorithms trained by the cuboids’ viability data, to more directly evaluate novel therapeutics or combinations in intact-TME human samples for the last stages of drug development.^50^ Although the heterogeneity (even of the baseline viability) can make the assay “noisy,” this variation in the drug responses likely reflects the differences in each cuboid, a strength of this approach. In a traditional bulk measure, a 100% response in half of the tissue gives the same instrument reading as a 50% response over the whole tissue. In our cuboid approach, a 100% response in half the cuboids looks very different from a 50% response in all the cuboids, allowing us to identify (and potentially further analyze) which part of the tissue corresponds to which response. Since our assay is non-destructive, downstream analysis of the cuboids, *e*.*g*., by RNAseq or proteomics, could provide further insights into the variability and underlying biological processes for cancer drug treatments.^50^

## Conclusions

Starting with a small biopsy microdissected into hundreds or thousands of TME-intact microtissues, our low-cost robotic workflow can reliably sort the microtissues into standard multi-well plates in less than one hour and gather a large amount of functional information such as drug efficacy data from fluorescent readouts, but different microtissue arrays with other outputs are also imaginable.^68,69^ Since the microfluidic mechanism for “lifting” microtissues can be accurately explained by theoretical fluid mechanics analysis, and it is insensitive to key parameters, it should be generally applicable to a variety of microtissue formats. Our data collectively shows the potential for our user-friendly robotic platform to allow minimally-trained biomedical researchers to conduct low-cost drug evaluations with live microtissues. We show that the platform can work with a range of microtissue sizes (250 – 400 μm), shapes (cuboidal and spheroidal), and species (mouse and human). We focused our study on microdissected tumor “cuboids” with intact TME^50^ because cancer drug evaluations suffer the most failures during the FDA approval process (more than 96%),^2^ so oncology is the area that can benefit the most from improvements. Our drug evaluation workflow with human cuboids highlighted the value of the exercise by revealing the response variability and small differences between tumor cores. This “heterogeneity information” can only be revealed with a multiplexed functional test such as ours and could in principle be used to inform clinical decisions, *e*.*g*., with a similar assay performed on cuboids from a biopsy of a patient who has never undergone chemotherapy yet. Hence, we believe that our platform could become an indispensable tool for the “democratization” of microtissue-based testing in a variety of scenarios, from drug development and disease models to personalized medicine.

## Materials and Methods

### Fabrication and operation of the rotary pump

The pump utilizes a rollerless eccentric design to generate peristalsis along a 1.3 ID silicone tube (See Rotary Pump in Suppl. Mater. for more details). The pump housing was CNC-milled with a 3-axis CNC mill (DATRON neo, Germany) on poly(methyl methacrylate) (PMMA) sheets of 8 mm thickness (McMaster-Carr, Elmhurst, IL). A central axle was 3D printed in polylactic acid (PLA, 3D Solutech), using a Flashforge Creator Pro™ Fused Deposition Modeling (FDM) 3D printer. The axle holds 3 metal Boca™ ball bearings of varying sizes, with the central bearing applying low friction peristalsis to the silicone tube, measuring 27 mm outer diam. (O.D.), 20 mm inner diam. (I.D.) and 4 mm thick. Two identical flanged bearings hold the central axle in place, measuring 18 mm O.D., 12 mm I.D. with a 0.8 mm-thick flange of 19.5 mm O.D. The flanged bearings provide the structural rigidity of the central axle to allow for low friction and controlled incremental movements using the stepper motor. The middle bearing is mounted eccentrically to the round inner PMMA casing. Four aluminum screws of 3 mm diameter lock the pump to the stepper motor on the end-effector of the robotic arm. Tube replacement can be easily achieved by removing the screws and opening the housing. Peristalsis occurs as the eccentric ring bearing rolls around the silicone tube, pressed against the inner housing. The peristaltic movement generates fluid (in our case, air) displacement in the silicone tube, causing liquid to be suctioned upwards into a glass capillary attached to the silicone tube, or dispensed downward out of the capillary, depending on the direction of rotation of the motor (See Rotary Pump in Suppl. Mater. for more details). When the pump rotates a full circle, the pumping rate is not constant due to asymmetries in the design of the pump and due to the presence of the inlet and the outlet. The angle origin (0°) is half-way between the pump inlet and the outlet. We observed a peak in the pumping rate at a 30 degree-wide region (from 180° to 210°) opposed to the inlet and outlet, so we typically operate the pump in that region for cuboid lifting (see **Fig. 2**B, left inset). The settings for the motor’s angular speed ω are in percent of maximum speed, so we had to calibrate the pump’s volumetric output as shown in **Fig. 2**B. Settings of 10% (0.5 rad/s), 25% (1.1 rad/s), 50% (1.4 rad/s), and 100% (1.8 rad/s) of the maximum ω produced ∼10.4 ± 2.2 μL/s, 24.6 ± 1.6 μL/s, 33.5 ± 1.9 μL/s, and 56 ± 6.1 μL/s of peak flow rate, respectively (blue curve in right inset of **Fig. 2**B). We were able to lift cuboids at 100% success with peak flow rates as low as 5.1 ± 0.3 μL/s (corresponding to a ω setting of 3%) and occasionally lifted cuboids with the pump’s ω set at 1% (but the lifting success diminished to 30%). In this work, the pump was always operated at 100% angular speed, resulting in a peak flow rate of 56 ± 6.1 μL/s and a total volume lifted of 6.4 ± 0.1 μL (cyan-shaded area in right inset of **Fig. 2**B).

### Cell culture

The Py8119 syngeneic mouse breast adenocarcinoma cell line (American Type Culture Collection (ATCC), CRL 3278) and U87-MG (ATCC) were grown in Dulbecco’s Modified Eagle Medium (DMEM)/F12 supplemented with 5% fetal bovine serum and 1% penicillin-streptomycin. Tissue culture reagents were obtained from GIBCO, ATCC, or Fisher.

### Tumor generation for mouse model

Mice were handled in accordance with institutional guidelines and under protocols approved by the Animal Care and Use Committee at the University of Washington, Seattle and by the Fred Hutchinson Cancer Research Center. For the Py8119 mouse syngeneic tumors, we injected 1-2 × 10^6^ cells in Matrigel (Corning) orthotopically into the mammary fat pad of >6-week-old female C57BL mice (Jackson Laboratories). For U87-MG human glioma cells xenograft tumors, we injected 1-2 × 10^6^ cells subcutaneously in 6-8-week-old male athymic nude mice (Jackson Laboratories). Tumors were harvested at <2 cm^3^. If not used immediately, the tumor was stored at 4 °C up to overnight in Belzer-UW cold storage medium (Bridge-to-Life Ltd).

### Human tissue

Human tissue was obtained with written informed consent and treated in accordance with Institutional Review Board approved protocols at the University of Washington, Seattle. The biopsy was from a 54-year-old male with recurrent colorectal cancer metastatic to the liver and peritoneum. He had previously received surgery, treatment with IR liver ablation, and chemotherapy with cisplatin, irinotecan, leucovorin, and 5-fluorouracil (5-FU).

### Cuboid generation and culture

We generated cuboids as previously described.^45^ For clarity, here mouse tumor cuboids are termed “spheroidal cuboids,” as they evolve into a round shape when cultured for a few days. We embedded tissue punches (600 μm diameter, Harris Uni-Core) in 1-2% lo-melt agarose and then cut slices using a Leica VT 1200 S vibrating microtome or MZ5100 vibratome (Lafayette Instruments). We cut the slices into cuboids with a tissue chopper (McIlwain tissue chopper (Ted Pella, Inc.), then gently dissociated the cuboids with flow using a transfer pipette filled with serum-free medium (DMEM/F12). For 400 μm cuboids, we filtered them with a 750 μm filter to remove oversized pieces and next with a 300 μm filter to remove smaller pieces (Pluriselect). Tissue was handled using ice-cold DMEM/F12.

After transfer of the cuboids to a 100 μm cell strainer (Corning or Falcon), we washed them twice with sterile PBS and once with medium. For the human CRC tumor, we did these washes in sterile tubes instead. For the CRC cuboids, the culture medium was Williams’ Media E (Sigma) supplemented with nicotinamide (12 mM), L-ascorbic acid 2-phosphate (50 mg/mL), D-(+)-glucose (5 mg/mL) from Sigma; sodium bicarbonate (2.5%), HEPES (20 mM), sodium pyruvate (1 mM), Glutamax (1%), and penicillin-streptomycin (0.4%) from Gibco; and ITS + Premix (1%) and human EGF (20 ng/mL) from BD Biosciences. For the mouse cuboid experiments, the culture medium was DMEM/F12 with 5% heat-inactivated fetal bovine serum and 0.1% penicillin-streptomycin.

### Robotic platform setup

To become operational, the robotic platform needs to go through a physical setup process (platform assembly and capillary alignment) and a calibration between the camera (ELP 4K IMX317 5-50 mm focus USB Camera) and the robotic arm. The robotic arm (DOBOT MG-400) is outfitted with a laser pointer for calibration (used as a reference point), the custom rotary pump connected to a glass capillary, and a custom workspace platform for housing the culture dishes and well plates. The camera is positioned with a holder above the microtissue-containing culture dish. To increase the contrast (tumor microtissues are opaque), the culture dish is back-illuminated by a flat white LED panel at all times. All devices that require power are plugged into a wall socket; the camera and robotic arm are connected to a PC with respective cables for data transfer. After the physical setup, a mapping between the robot’s and the camera’s system of coordinates needs to be established. The mapping is established with the assistance of a special calibration dish that is placed in the field of view of the camera. The robotic arm moves through the predefined positions located on the calibration dish, while the camera records the position of the laser dot in the calibration dish. Once the positions have been recorded, a coordinate transformation matrix can be calculated, which finishes the calibration process (see Computer Vision in Suppl. Mater., and **Fig. S3**A&B). The capillary is aligned to the reference point of the robotic arm using a custom dish with a conical hole drilled in it (see Capillary Alignment in Suppl. Mater.). The robotic platform is then ready for operation. To run the picking procedure, the user places a random distribution of microtissues (cuboids or spheroidal cuboids) in the culture dish. The software takes a picture of the culture dish and registers the coordinates of each microtissue’s projected (2D) center of mass. These coordinates are used to direct the capillary to particular microtissues. Before starting, the user is prompted to specify the range of microtissue sizes to pick from and parameters to ensure selection of isolated microtissues. The user selects the number of microtissues to transfer and defines the destination well-plate format (or any other array of positions).

### Programming of the automatic tissue transfer procedure (GUI)

The user is required to initialize several input variables that affect the automatic tissue transfer procedure through a Graphic User Interface (GUI) freely available at https://github.com/steiva/cuboid-manipulator-distribution. These variables include the picking size range, minimum distance to nearest neighbor, indexes of the wells to fill (in other words, how many cuboids to transfer), etc. The user must also choose the target type (a 96- or 384-well plate or a set of locations). The transfer procedure can then be started. Images taken by the camera are analyzed by the software and information on the location of the microtissues is given to the robot. The software analyzes the distribution of the microtissues and filters out those that do not fit the preset conditions. With activation of the rotary pump, the robotic arm suctions ∼6.4 ± 0.1 μL of fluid, which lifts an automatically selected microtissue and reverses the pumping direction to deliver it to the target well location. In the unlikely event that microtissues within the culture dish do not satisfy any picking conditions, the process auto-pauses so their distribution can be reset by the user, here by manually shaking the culture dish. The robotic arm then continues the transfer procedure until all 96 or 384 transfers to the well plate have occurred, at which point the procedure will end. During the transfer process, the software can determine failed transfer attempts, then rectify them by refilling wells that were left empty unintentionally. Using this straightforward protocol, the robotic platform can fill a 384-well plate with single microtissues per well in ∼40 min. Using a protocol that loads multiple cuboids at once into the capillary, the platform could potentially fill a 384-well plate with up to 6 cuboids per well in ∼133 min.

### Drug treatment and imaging

To measure the viability of human tumor cuboids, RealTime-Glo (Promega) was added at 1x or 0.5x and the baseline luminescence was read the following day by IVIS (Perkin-Elmer). Drug was added to the well, and luminescence was read again after incubation with drug without further addition of RealTime-Glo. To measure cell death in mouse tumors as an endpoint, SYTOX Green (1/50,000; Invitrogen), and/or Hoechst (16 μM, Invitrogen) were added to the well, incubated for 1 hr at 37 °C, then imaged with or without washing twice in PBS. We took bright-field images using a Canon DS126601 camera on a Nikon SMZ1000 dissecting scope and fluorescence images using a Keyence BZ-X800 microscope. Graphing and statistics were performed on GraphPad Prism, with power analysis performed on clincalc.com. Drugs were purchased at MedChem Express except for staurosporine (Thermo Scientific) and for fluorouracil, oxaliplatin, irinotecan, regorafenib, and fruquintinib (Selleck).

### Data acquisition, processing, and visualization for Py8119 mouse tumor

We imaged the plate under blue and green filters using a Keyence BZ-X800 microscope to locate the microtissues as well as their respective average cell death fluorescence signal from the green channel image. We used the blue channel (Hoechst) to determine the perimeter of the microtissues. We measured the average green channel fluorescence within each perimeter using ImageJ. We subtracted the background fluorescence (green channel) from each cuboid and sorted all values in Microsoft Excel. Histograms were then created in Prism displaying the average green channel fluorescence (representative of cell death) for each drug condition, with each individual cuboid represented by data points over their respective bar graph. The standard error of the mean is displayed and the significant differences between the conditions and the control groups are noted with asterisks.

### Statistical Analysis

GraphPad Prism 9 was used for tests of significance which are done as indicated. Post-hoc power analysis was performed using https://clincalc.com/stats/samplesize.aspx.

## Supporting information

Supplementary materials full document

## Acknowledgments

We thank Heidi Kenerson for preparing the human tumor tissue slices.

## Author contributions

Conceptualization: AF

Methodology: AF, LFH

Investigation: IS, NRG, EJL, LFH, TNHN

Formal Analysis: AA, LFH, IS, NRG

Visualization: IS, NRG, LFH

Software: IS, DS, SH

Data Curation: LFH, IS, NRG

Supervision: AF, LFH, TSG, RSY

Writing—original draft: AF, IS, NRG, LFH

Writing—review & editing: AF, LFH, IS, NRG

## Data and materials availability

All data are available in the main text or the supplementary materials with the exception of original well-plate images, which are available upon request.

## Supplementary Materials

Fig. S1. Setup of the robotic platform.

Fig. S2. Robotic platform within a sterile tissue culture hood highlighting individual components.

Fig. S3. Computer vision.

Fig. S4. Graphs of transfer success for a total of 6 experiments.

Fig. S5. Combined fluorescence readout of drug responses for two 96-well plates filled simultaneously.

Fig. S6. Design of the rollerless eccentric pump.

Table S1. Existing commercial robotic platforms.

Table S2. Size selection.

Table S3. Microtissue transfer success statistics for mouse and human tissue tumors.

Movie S1. Demonstration of cuboid localization and suction by the capillary.

Movie S2. Demonstration of the operation of the custom peristaltic pump.

Movie S3. Close-up of cuboid lifting.

Movie S4. Demonstration of the operation of the robotic platform.

Data S1. (separate file) Rollerless eccentric peristaltic pump design file (.dwg).

Data S2. (separate file) Rollerless eccentric peristaltic pump design and assembly descriptions (.pdf).

