## Supplementary materials full document for "Low-Cost Robotic Manipulation of Live Microtissues for Cancer Drug Testing"

Ivan Stepanov et al.

This PDF file includes:

Supplementary Text
Figs. S1 to S6
Tables S1 to S3

Other Supplementary Materials for this manuscript include the following:
Movies S1 to S4
Data S1 to S2

### Supplementary Text

#### Rotary Pump

Design background

A rollerless eccentric pump was fabricated to allow smooth, non-pulsatile fluid transfer as compared to traditional tri-roller peristaltic pumps (**Fig. S6**). The ring moves against the tube in one radial motion, only losing tube contact once per revolution, yielding fewer pulses than typical tri-roller peristaltic pumps (**Fig. S6**A&B)**.** The pump is attached to the robot stepper motor to allow for direct continuity with the robot API (**Fig. S6**A). A minimum suction volume of ~6.4 ± 0.1 µL within the capillary is required to "lift" and transfer individual microtissues from a culture dish into a well plate. The stepper motor is limited to rotations between -360° and 360°, thus microtissue suction and deposit must be accomplished under such restrictions. The pump is designed to move ~30 ± 2 µL of fluid per revolution of the stepper motor to allow for flexibility in microtissue suction and delivery volumes. Smooth movement increments were considered in the design to minimize stress on the stepper motor and provide small rotation increments, only limited by the stepper motor incremental range of 0.1-degree movements.

Design process

A flexible silicone tube of appropriate volume and diameter was chosen in combination with a rigid, transparent 6.5 mm thick PMMA (polymethyl methacrylate) housing. **Fig. S6**C shows close-up of housing and blue liquid within silicone tube. A PLA central axle was designed to hold the ring-ball-bearing eccentrically to the housing to cause peristalsis from movements of the stepper motor. Two additional flanged-ball-bearings were used to hold the central ring bearing within the PMMA housing. The three bearings (two: MF6701-ZZC PS2, one: SMR6704-ZZC #3 PS2) were ordered from Boca Bearings. All additional components were fully modeled in Autodesk Fusion 360 CAD software.

Fabrication

The pump CAD design (**Fig. S6**B&D) was converted from STL format to x3g format using the FlashPrint slicer tool. Fabrication consisted of 3D printing the central axle (where the ring bearing is rigidly attached in the middle) with PLA filament using a FlashForge Creator Pro FDM 3D printer. Two flanged bearings hold the central axle in the middle of the two PMMA housing pieces. The central axle contains an asymmetric disc in the center where the ring-ball-bearing is pressure fitted. Upon completion of the CAD model, the housing pieces were translated to ISO G-code using Autodesk Fusion 360 for CNC milling. Milling was a ~2-hour-long procedure; a 12.5 mm thick PMMA square (30.48 x 30.48 cm, McMaster-Carr) was milled into the desired housing parts using a Datron Neo 3-axis CNC milling machine. The pump was then assembled with all seven parts: top housing, bottom housing, silicone tube, two flanged bearings, large ring bearing and 3D printed central axle. The housing contained 4 female-male post connectors to rigidly interlock the top and bottom pieces. Four hex screws were used to fasten the pump to existing threaded openings on the stepper motor mount. Adequate tube length was used to accommodate the housing and allow room for attachment of a glass capillary tube used as the end-effector. Detailed design drawings are provided in "Suppl. Data S2 pdf". See "Data S1.dwg" for access to the CAD file. The following link is also provided for the fusion 360 file: https://a360.co/3RXI08s

### Capillary alignment

The capillary needs to be aligned to the laser pointer to make the robotic platform able to localize and suction microtissues. The accuracy with which the microtissues are picked up depends on the capillary alignment, which requires trial and error for a better result. To facilitate the alignment process, a special culture dish was made. The culture dish has a small hole drilled in the center, with a diameter smaller than that of the capillary. The hole is then widened at the surface of the culture dish, forming a funnel. The funnel is colored black, so that it is easier for the software to detect. The robot is instructed to move to the position of the funnel using the software. The circle is now acting as a reference point for the location of the laser, making it easier for the user to complete the physical alignment of the capillary to the laser pointer. The capillary is positioned to fit into the funnel, touching the culture dish so that the circle of the capillary would match with the reference hole. The capillary is now aligned with the laser and can be used to transfer tissue.

### Computer vision

#### Robot calibration

The camera's system of coordinates is represented by a discrete two-dimensional cartesian system consisting of rows and columns of pixels, whereas the robot's system is a three-dimensional cartesian system, measured in millimeters. When the robot and camera are first initialized, there is no mapping between the two coordinate systems. The robot calibration procedure creates a transformation matrix that maps pixel coordinates into robot coordinates.

To obtain the transformation matrix, we need to record a series of positions of the robot, while recording visual marks related to the robot's movement. The marks are created by a laser that is physically attached to the robot. When the robot moves to one of the calibration positions, the laser creates a bright spot, called an anchor, on the working platform that is then detected on an image taken by the camera. A series of coordinate pairs is obtained, where each coordinate of the robot is tied to the corresponding pixel coordinate of an anchor. The details of this process are discussed below.

##### *Anchor region masking*

The calibration benefits from finer visual marks. When the laser shines on a white surface, the spot it creates is visually larger than when it is shined on a darker surface due to more light reflection. Hence, black anchor regions are created for the laser. These anchor regions are easy to detect (see **Fig. S3**A) using conventional computer vision methods. Binary thresholding is applied to a grayscale image, followed by opening and erosion morphological operations. A binary mask is then created with the resulting contours.

The mask can now be applied during the actual calibration process to increase its robustness. A reset of the mask is only required when the physical setup changes and/or anchor regions are shifted, if the setup does not change throughout usage, the creation of the mask can be done only once.

*Recording of the calibration coordinate pairs*

Coordinate pairs need to be recorded to establish a mapping between the robot and the camera. The robot travels to 4 pre-defined positions on a single height and is being polled for its coordinates at each location, which are then recorded (see **Fig. S3**B). The camera takes a snapshot at each of the 4 positions and each image is analyzed to find the anchors. At each position the laser point is detected by using thresholding and attributed to the corresponding spatial coordinate of the robotic arm. We obtain a data structure of the following form:

$$\begin{Bmatrix} (rX_1, rY_1) : (cX_1, cY_1) \\ (rX_2, rY_2) : (cX_2, cY_2) \\ (rX_3, rY_3) : (cX_3, cY_3) \\ (rX_4, rY_4) : (cX_4, cY_4) \end{Bmatrix}$$

Here, *rX* and *rY* represent the robot coordinates and *cX* and *cY* the corresponding pixel coordinates. We are now ready to compute the transformation matrix.

*Transformation matrix calculation*

The features of interest lie in a single plane (microtissues in a culture dish). Hence, a linear relationship can be established between the image plane and the plane of interest in the 3D world. Based on the pinhole camera model, the relation between $r = [rX, rY, 1]$ and $c = [cX, cY, 1]$ can be expressed as $r = Tc$, where $T$ is a transformation matrix.

$$r = Tc \rightarrow \begin{bmatrix} rX_i \\ rY_i \\ 1 \end{bmatrix} = \begin{bmatrix} t_{11} & t_{12} & t_{13} \\ t_{21} & t_{22} & t_{23} \\ 0 & 0 & 1 \end{bmatrix} \begin{bmatrix} cX_i \\ cY_i \\ 1 \end{bmatrix}$$

Reorganizing the equation, we get:

$$\begin{bmatrix} cX_i & cY_i & 1 & 0 & 0 & 0 \\ 0 & 0 & 0 & cX_i & cY_i & 1 \end{bmatrix} \begin{bmatrix} t_{11} \\ t_{12} \\ t_{13} \\ t_{21} \\ t_{22} \\ t_{23} \end{bmatrix} = \begin{bmatrix} rX_i \\ rY_i \end{bmatrix}$$

To solve this system of equations, we need at least 3 coordinate pairs. With the least squares method, we can solve an overdetermined system of equations. In our implementation, 4 coordinate pairs are sampled in the camera-to-robot calibration process. Solving the system of equations, we obtain the transformation matrix, which can be applied to the camera coordinates as in the previous equation to obtain the robot coordinates.

*Well plate grid calibration*

The coordinates of wells in the well plate are calculated based on its four corners.

The robot is used to record the coordinates of the four corner wells of the well plate. The capillary must be aligned beforehand, as the coordinates are being recorded with its tip. Having 4 pairs of X, Y coordinates, we can divide the intermediate space between them into a grid based on the size of the well plate. The coordinate grid is then recorded to a file and can be used in the future.

Microtissue recognition

We calculate the positions of the microtissues within a culture dish based on simple image processing methods combined into a conventional Computer Vision pipeline. The culture dish where the tissues reside is located first by a circle detection algorithm. Binary thresholding is applied to the area inside the culture dish to locate the tissue samples. Thus, a list of microtissue contours is obtained. Prior to picking the microtissues, the contours need to be analyzed. We perform the following steps during the analysis:

1. *Derivation of the microtissue center coordinates* from the contour information (giving every microtissue a distinct coordinate position).
2. *Prevention of picking non-tissue material*, such as debris that is possibly present in the target Petri Dish. The debris can be filtered out based on its size – the area of the contours of the debris is commonly smaller than that of actual microtissues. Filtering can be done easily once the contour area is calculated (see **Fig. S3**D).
3. *Prevention of detecting bubbles as microtissues*. Occasional bubbles present in the culture dish can cause an empty well but the software deals with them preventively. If unchecked, the software would detect the bubble as a microtissue and attempt to transfer it but generate an empty well. To ensure bubbles are never picked up, the software checks if the microtissue's inner pixels are brighter than the edge pixels. If so, the microtissue is removed from the picking queue and classified as a bubble. However, if the bubble is small, there may not be enough resolution to determine the brightness of its center. In the case where the target size to be picked is on the smaller range (~250 µm), the platform can recognize as microtissues and attempt to pick up smaller bubbles

that would otherwise be filtered out based on size, resulting in an empty well (see **Fig. S3**D).

4. *Prevention of acquiring more than one microtissue at a time*. Acquiring more than one microtissue at a time can happen when a microtissue is too close to the target one. The determination of the most isolated microtissues is done by building a 2-D binary tree where nodes are coordinated pairs of the detected microtissues. The 2-D binary tree allows us to find nearest neighbors for each node, enabling us to find the most isolated microtissue. A minimum distance threshold which needs to be surpassed for microtissues to be picked is set.

<u>Size selection configuration</u>

The robotic platform enables finely-tuned tissue size selection for transfer. We evaluated camera noise by taking 1,000 micrographs of a single cuboid and using the Microtissue Recognition tool (see above) for each image, which resulted in an average cuboid size of 412 µm ± 3.52 µm (max. and min. recorded sizes: 422 and 404 µm, respectively). A series of experiments was conducted to quantify the software's size selection accuracy and characteristics. A culture dish was filled with ~500 cuboids, and upon acquisition of 100 images an average size histogram was created by averaging the size histograms (displaying the size of each cuboid within the culture dish) gathered for each image. The average histogram was then normalized to obtain the size frequency (see Microtissue size distribution analysis for detailed characteristics of the size selection feature). A picking size range is then selected centered around the middle of the normal distribution of tissue size (the highest peak). The robot is programmed to transfer 100 tissue samples within the defined size range to an empty culture dish. After the transfer is complete, the size distribution of the 100 tissue samples in the target culture dish is analyzed. The results are saved, and the microtissues in the target dish are transferred back to the source dish. The same procedure is conducted two more times, with new initial distributions and with decreased picking size windows (see **Fig. 3**A). Two more series of the same three experiments are conducted to sample the repeatability of the results.

*Size conversion ratio*

Area size (in $\mu m^2$) of microtissues is acquired using a conversion from pixel area to metric area.

*Pixel area to square microns conversion*

Microtissue contours are recorded upon detection on an image. The area of the contour is calculated with a built-in function in OpenCV-python. The function returns a float value employing Green's theorem to calculate the area of an enclosed contour. The value does not reflect the actual physical size of a microtissue, as the pixel density per microtissue changes with any alterations of the physical setup.

Conversion is based on a simple ratio between the tissue sample and an object of known size. In the case of our project, a triangular object was manufactured out of black plastic using a laser cutter. The object was measured using a Keyence VHX-7000 series microscope.

We detect the triangle in an image and record its area in pixels (see **Fig. S3**C). We apply inverse binary thresholding to the image, followed by a dilation operation. The contours on the resulting image are filtered by their shape to find the triangular object, and the pixel area of the triangle is calculated.

Having found the area of the reference triangle, we can now proceed to calculating the ratio:

$$R = \frac{S_{ref}[\mu m^2]}{S_{ref}[pixels]} = \frac{S_{cuboid}[\mu m^2]}{S_{cuboid}[pixels]}$$

where $S_{ref}$ is the area of the reference object, and $S_{cuboid}$ is the area of a microtissue. The unknown in this equation is $S_{cuboid}[\mu m^2]$, and having $S_{cuboid}[pixels]$, the calculation is trivial:

$$S_{cuboid}[\mu m^2] = R \cdot S_{cuboid}[pixels]$$

We are now able to calculate the microtissue area in micrometers squared, or, by extension, any metric size unit.

*Conversion from microtissue area to its effective diameter*

The *effective diameter* of the microtissue is calculated. We assume the microtissue projection on to the image plane to be circular, hence its diameter is derived as follows:

$$S = \pi r^2 = \pi \frac{d^2}{4} \Rightarrow d = 2\sqrt{\frac{S}{\pi}}$$

The effective diameter is then used as the main metric for size selection during the microtissue manipulation by the robot.

*Microtissue size distribution analysis*

We obtained the size distribution of the microtissues in the culture dish from an aggregate histogram produced from 100 images (to reduce sensor noise).

Automated picking procedure

The automatic picking procedure is used for filling up well plates with microtissues. The picking procedure starts by analyzing the culture dish with microtissues. If there are microtissues that pass all the preventative checks, they are marked as “pickable”. The robot then proceeds to picking a random “pickable” microtissue from the distribution and carrying it to the destination well plate.

**Transfer Success**

The calibration of the robotic platform as well as the manual capillary alignment introduce a compounded error into the robot's ability to localize a target microtissue sample. To operate the robotic platform in an informed manner, we conducted a study of the tolerance for compounded error. We put a single microtissue into a culture dish and recorded its size and shape. The culture dish was filled with 4 mL of 1% BSA in PBS. The robot was given instructions to pick a single tissue sample in a grid of spatial positions ((z, r) parameter space, increments of 0.1 mm) from z = 0.5 mm above the culture dish and r = 0.0 mm radial offset from the microtissue in the XY plane. For each spatial position, picking was attempted 10 times, with the pickup success being recorded. The result was a heatmap of the pickup success rate, which gave insight into the robot's accuracy in localizing the microtissue while still being able to pick it up.

The experiment was conducted a total of 6 times: Experiment group A - 3 times on 3 different microtissues (368 µm, 395 µm and 360 µm effective diameter) and experiment group B - 3 times on a single 314 µm spheroidal cuboid. All experiments were conducted with different calibrations of the robotic platform (**Fig. S4**A&B). In one of the group B experiments, the heatmap was shifted down by 0.2 mm. The shift was due to human error: the capillary was not fully pressed down into the culture dish during capillary alignment, producing this offset in the experimental results. Three important conclusions can be made from these experiments. Firstly, the system is reliable regardless of any random human error made while aligning the capillary – the height at which the microtissue is picked with 100% success ranges from $z$ = 0.5 mm to $z$ = 1.0 mm. Secondly, we can make an informed decision on how to set the minimum distance to nearest neighbor for picking microtissues, while avoiding suctioning two microtissues at the same time. Based on **Fig. S4**, we set this distance to be 2.0 mm (which is well into the safe zone). Lastly, the platform works reliably for both spheroidal- and cuboidal-shaped microtissues.

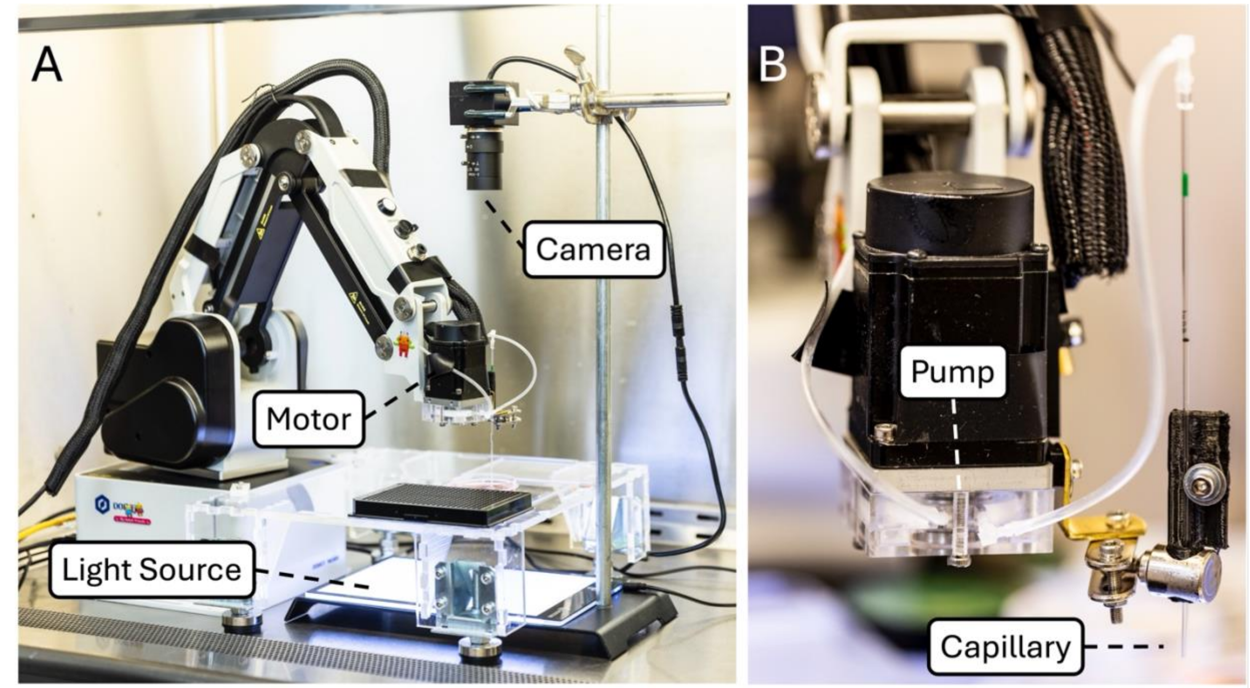


**Fig. S1. Setup of the robotic platform**

**(A)** The platform inside a tissue culture hood displaying the camera positioned above the culture dish and well plate, with the LED light source underneath. **(B)** Close-up of the robot's "head" illustrating the mounting of the pump and the capillary.

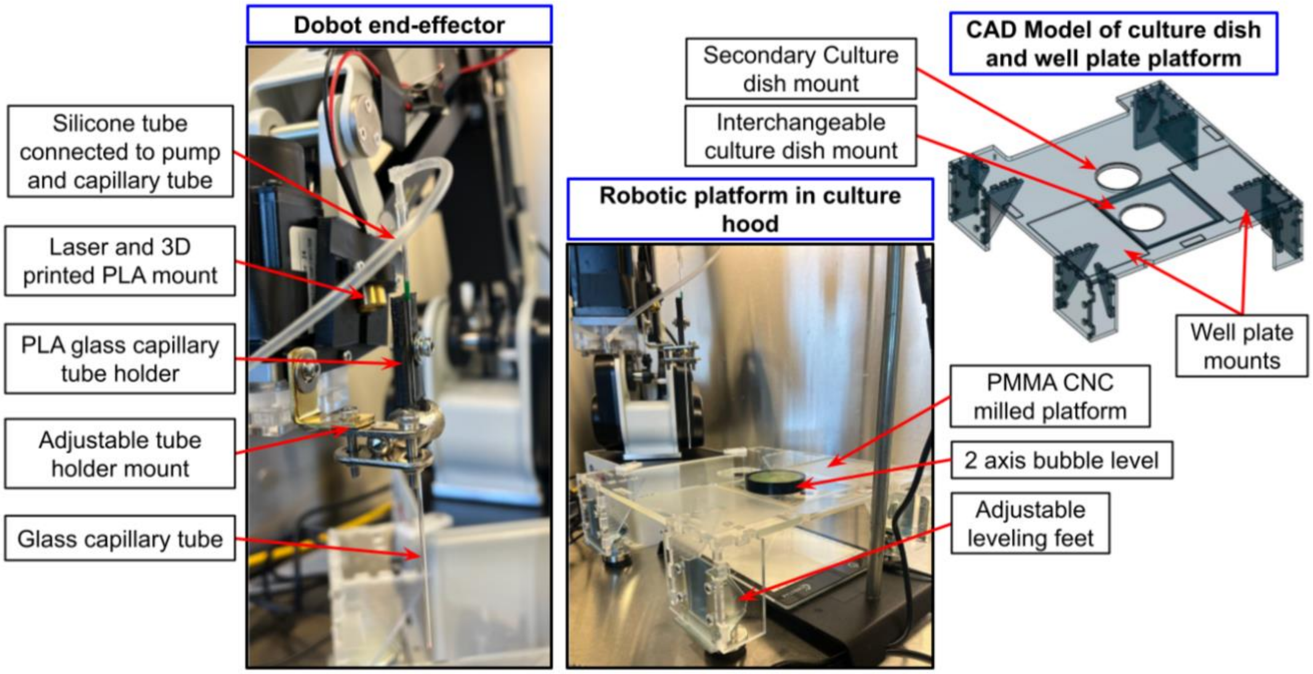


**Fig. S2. Robotic platform within a sterile tissue culture hood highlighting individual components**

The platform was designed in Fusion 360 CAD software to attach level to the accessible area of the DOBOT MG400 via two 3D-printed poly-lactic acid (PLA) brackets. PMMA of 6.5 mm thickness was cut to the desired shapes using the CAD model on a Datron Neo 3-axis CNC milling machine. Each piece was fitted together using jigsaw carpentry techniques, so no screws or glue were needed for assembly. Height adjustable metal leveling feet (Tahikem 4-pack heavy-duty levelling feet) were attached to the four legs of the PMMA platform with four screws each. Two interchangeable culture dish holders were milled on 2 mm-thick PMMA to allow for standard size usage of 60- and 100-mm diameter culture dishes. The laser mount and the glass capillary holder were 3D printed in PLA. A 635 nm-wavelength, 100 mm-focus laser diode (VLM-635-63 LPO-100, Quarton Inc.) was installed as the calibration laser for the platform. A voltage regulator (LM317MTG, Onsemi) was soldered to a 32 Ω (½ W) resistor (CMF5532R000BHEK, Vishay Dale) and then soldered to the laser and a USB-A connector for connection to the computer for power. Metal micromanipulator ball joints and box brackets were used to create an adjustable end-effector to hold the glass capillary holder. A 2-axis bubble level is used to measure the levelness of the platform which is adjusted with the leveling feet.

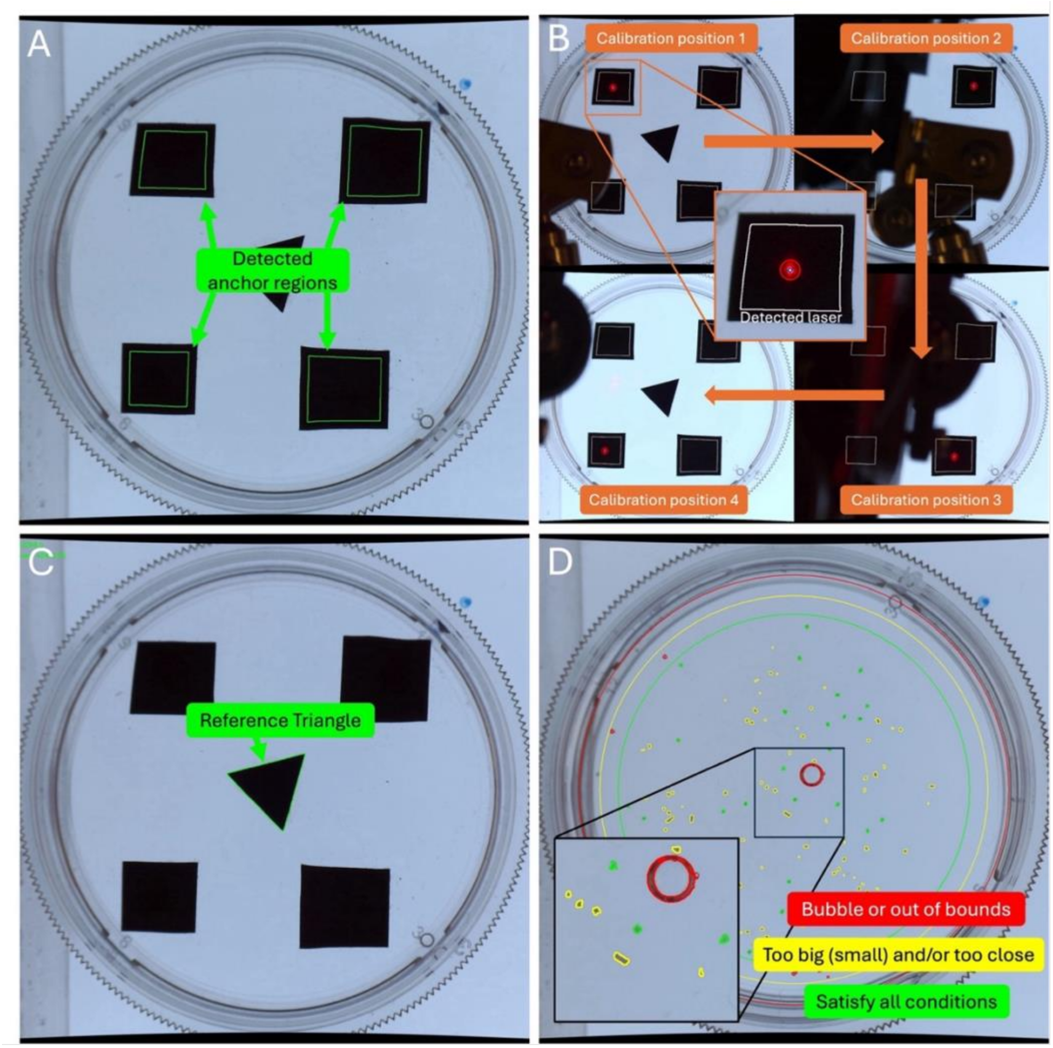


**Fig. S3. Computer vision**

**(A)** Anchor regions detected by the software (in green). A mask is made from these detected regions and applied during the calibration. **(B)** Visualization of the robot calibration process: The robot moves through calibration positions 1, 2, 3 and 4, recording its coordinates. At each position, the laser anchor is detected, and the pixel coordinates are recorded. The corresponding robot and pixel coordinates form a coordinate pair that is then used for calculating the transformation matrix. **(C)** Method for establishing a size conversion ratio using an object of known dimensions: The reference triangle detected on an image. This triangle has a known size and can be used to calculate a size conversion ratio. **(D)** Detected and filtered microtissue: Outer red ring: ring detected by the software to filter out everything outside the culture dish. Middle yellow ring: software starts detection of microtissues inside this ring, but never picks them before the inner green ring. Inner green ring: all microtissues are detected and can be picked by the robot. If an object is labeled in red, it is out of picking bounds or is detected as a bubble. If an object is labeled as yellow, it is a microtissue that does not satisfy size conditions and/or is too close to other microtissues. If a microtissue is labeled green, it satisfies all conditions and will be picked up by the robot.

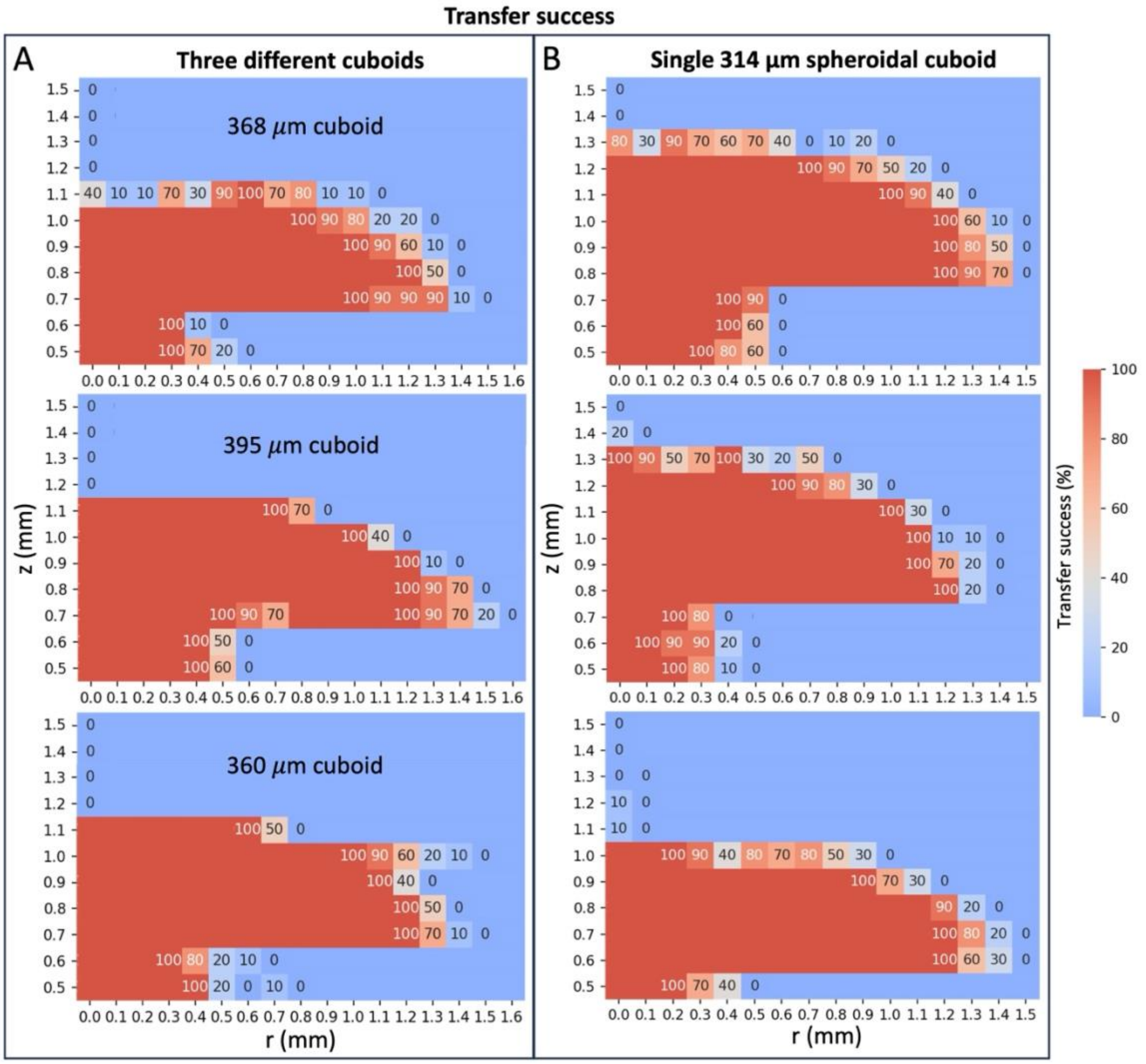


**Fig. S4. Graphs of transfer successes for a total of 6 experiments**

Transfer success graphs as a function of radial distance *r* from the cuboid's center and the height *z* of the capillary above the culture dish bottom, with color indicating transfer success percentage. Red zones are 100% transfer success and blue zones are 0% success. Experiments were conducted with different calibrations of the robotic platform. To make cuboids of two different shapes, we fixed mouse cuboids at Day 0 to obtain cuboidal microtissues and at Day 3 to obtain spheroidal microtissues. **(A)** Transfer success graphs for three different fixed PY8119 mouse breast cancer cuboidal microtissues: sizes 368 µm, 395 µm and 360 µm. **(B)** Three transfer success graphs for a single 314 µm Py8119 spheroidal tumor microtissue.

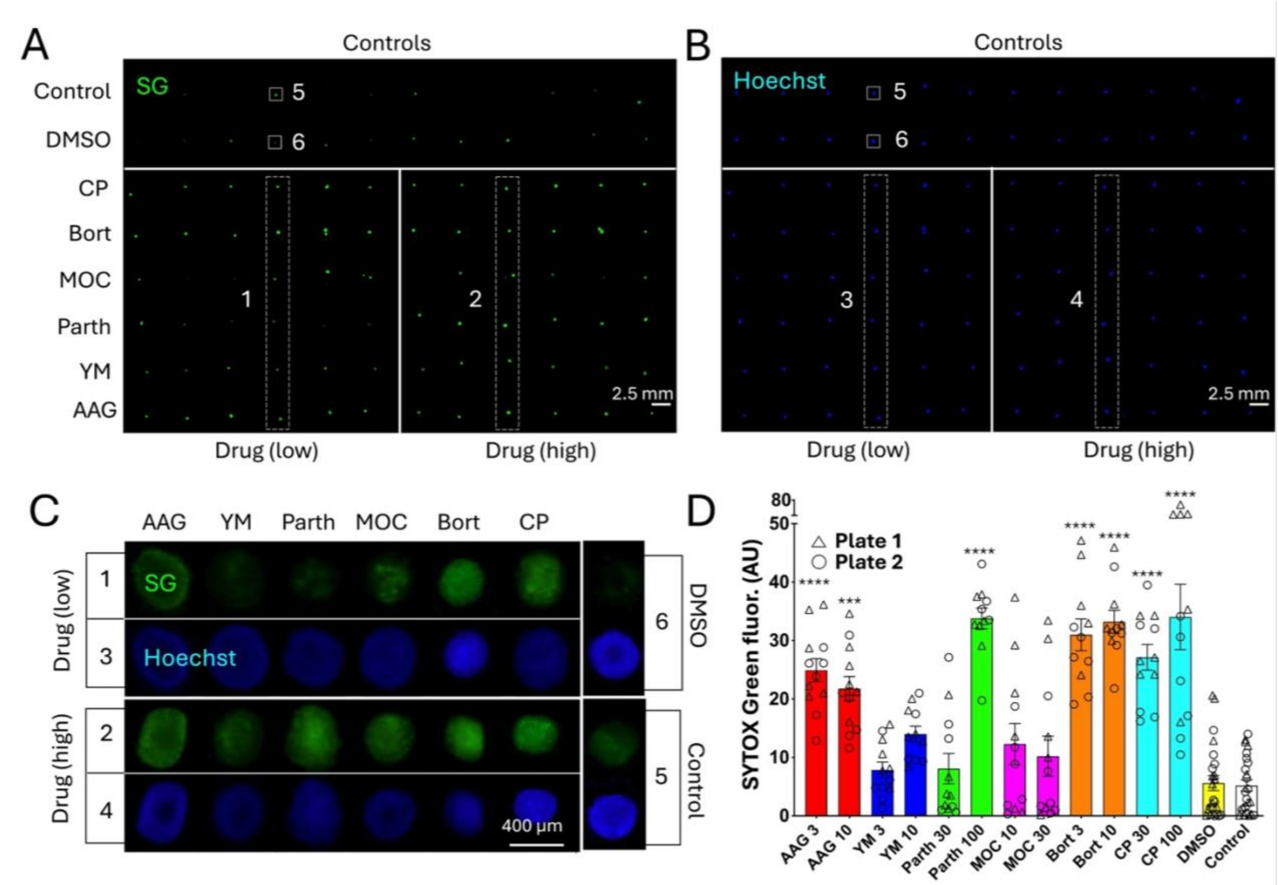


**Fig. S5. Combined fluorescence readout of drug responses for two 96-well plates filled simultaneously**

(**A&B**) Fluorescence images depicting 96 well plates loaded with U87 mouse tumor spheroidal cuboids. A total of ~192 cuboids (400 µm) were loaded individually into two 96-well plates and were then exposed to the indicated drugs in their respective concentrations (µM) for 3 days. Upon completing the incubation period (72 hrs), the cuboids were stained with the cell death indicator SYTOX Green (SG) (**A**) and Hoechst nuclear stain (**B**). **(C)** Close-up micrographs of both high and low drug concentrations on both green (SG) and blue (Hoechst) channels from plate 1**.** **(D)** Graph of the SG fluorescence intensities from both plates**.** Ave ± sem, each point represents a cuboid. n=12 wells per drug condition, n = 24 wells for control and DMSO Kruskal-Wallis test with Dunn post-hoc. versus DMSO except for CP (versus control) **** $p < 0.0001$, *** $p < 0.001$.

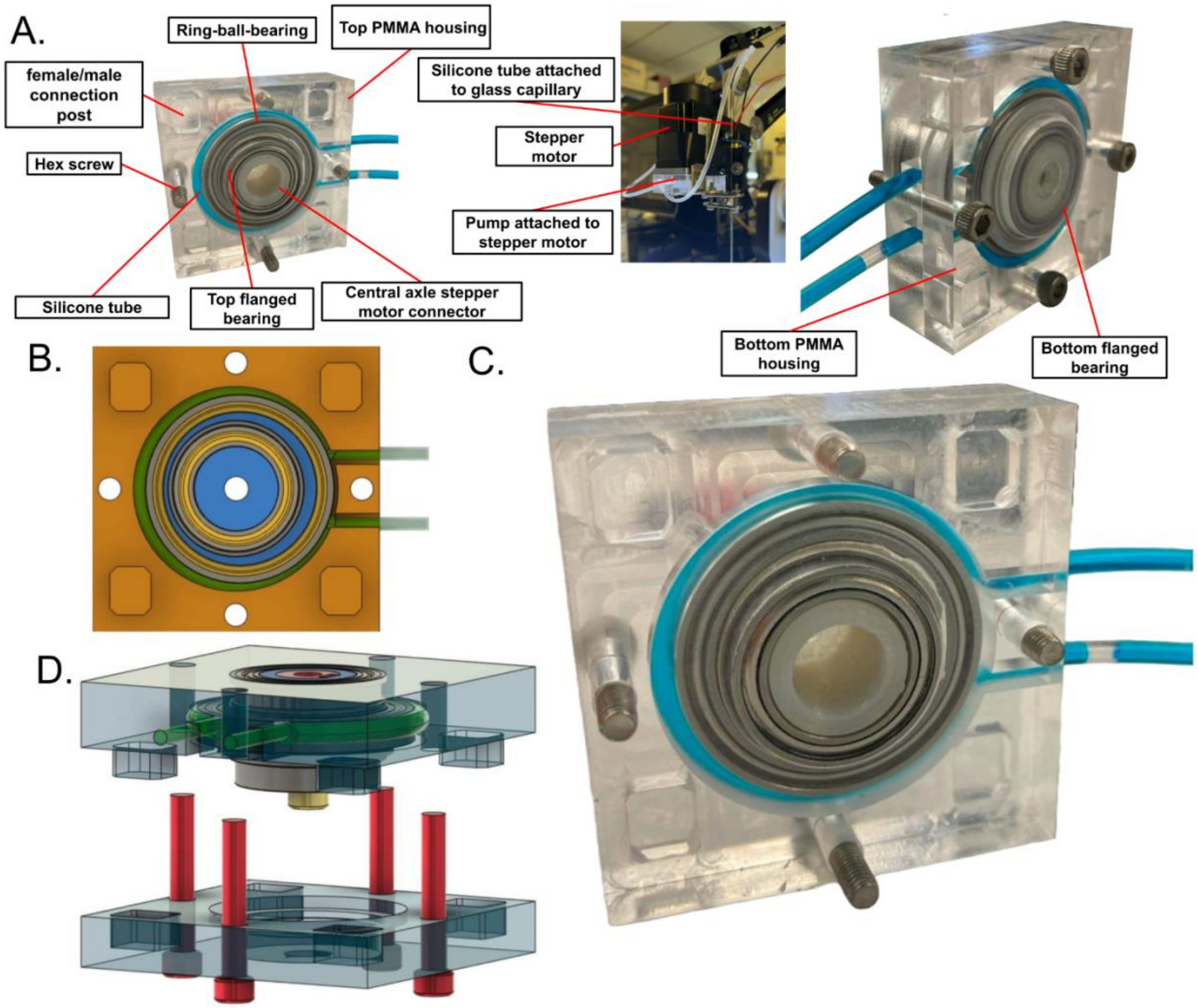


**Fig. S6. Design of the rollerless eccentric pump**

(**A**) Description of the parts used to fabricate the pump (left and right: bottom view; center: side view) and its attachment to the DOBOT MG400 stepper motor. (**B**) Top view of Fusion 360 CAD model used for the pump's design, illustrating the eccentricity of the central bearing in relation to the circular housing. (**C**) Close-up of the fabricated pump, containing blue-colored fluid within the tube for visualization purposes only. During real operation, the silicone tube is not filled with liquid and only displaces air. (**D**) Exploded 3D view of the pump's CAD model illustrating the assembly order. Please see Suppl. Data S2.pdf for detailed assembly, design and fabrication information.

**Table S1. Existing commercial robotic platforms.**

| Robotic Platform | Features | Loading time | Pricing |
|---|---|---|---|
| Molecular Devices ClonePix 2 | Transfer/select mammalian cells/ colonies.<br>Image analysis. | >10,000 mammalian colony clones selected and transferred per day | $325K - $650K |
| Molecular Devices QPIX | Transfer/select cells and cell colonies.<br>Image analysis.<br>Fluorescence.<br>Fluid handling. | 30 minutes per 96 well-plate. | $200K - $500K |
| Hudson Robotics RapidPick SP | Transfer/select bacterial colonies. | 25 minutes per 96 well-plate<br><br>Picks 250 colonies per hour | $83K |
| Cytena B.SIGHT | Bacterial colony/small organoid dispenser.<br>Image analysis.<br>Fluorescence. | 3 minutes per 96 well-plate | $180K - $210K |
| Shimadzu Cell Picker | Transfer/select individual cells and cell colonies. | N/A* | N/A |
| Sartorius CellCelector | Transfer/select cells and cell colonies.<br>Temperature control.<br>Fluorescence.<br>Image analysis. | 60 minutes per 96 well-plate. | $200K |
| Yamaha Cell Handler | Transfer/select organoids, spheroids and single cells.<br>Temperature control.<br>Fluorescence.<br>Image analysis.<br>Fluid handling. | 30 minutes per 96 well-plate. | $250K |

* Not available in the United States.

**Table S2. Size selection.**

| Range | Experiment 1 | | Experiment 2 | | Experiment 3 | | *Average enrichment %* |
|---|---|---|---|---|---|---|---|
| | Before % | After % | Before % | After % | Before % | After % | |
| **130 µm** | 82 | 95 | 81 | 97 | 84 | 89 | *11.3* |
| **90 µm** | 72 | 91 | 73 | 93 | 70 | 91 | *20* |
| **50 µm** | 47 | 82 | 49 | 81 | 47 | 72 | *30.7* |

**Table S3. Microtissue transfer success statistics for mouse and human tissue tumors.**

| Mouse tumor | | | | | | |
|---|---|---|---|---|---|---|
| **Type** | **Wells filled** | **Empty wells** | **Doubles** | **% Empty** | **% Double** | **% Success** |
| **Py8119** | 384 | 2 | 5 | 0.52 | 1.30 | 98.18 |
| **Py8119** | 384 | 2 | 3 | 0.52 | 0.78 | 98.70 |
| *Average* | | *2* | *4* | *0.52* | *1.04* | *98.44* |

| Human tumor<br>* *Note: some human tumors were too small to fill a whole 384-well plate*<br># CRC1 and CRC2 designate CRC Patients 1 and 2, respectively | | | | | | |
|---|---|---|---|---|---|---|
| **Type** | **Wells filled*** | **Empty wells** | **Doubles** | **% Empty** | **% Double** | **% Success** |
| **CRC1# Plate 1** | 342 | 24 | 16 | 7.02 | 4.68 | 88.30 |
| **CRC1 Plate 2** | 352 | 22 | 20 | 6.25 | 5.68 | 88.07 |
| **CRC1 Plate 3** | 352 | 6 | 5 | 1.70 | 1.42 | 96.88 |
| **CRC2#** | 384 | 9 | 10 | 2.34 | 2.60 | 95.06 |
| **ICC** | 384 | 15 | 14 | 4.26 | 3.98 | 91.76 |
| *Average* | | | | *4.31* | *3.67* | *92.01* |

**Movie S1. Demonstration of microtissue localization and suction by the capillary.**

(Left) The capillary hovers above the microtissue, lowers itself and locates the tissue sample. On the right pane, the accuracy and precision of the localization can be seen. The microtissue is then suctioned by the capillary and lifted above the culture dish. The microtissue is ejected back into the culture dish, thus showing the full cycle of the robot's operation: suction, transportation, and ejection of the microtissue.

**Movie S2. Demonstration of the operation of the custom peristaltic pump.**

The rotation of the pump from 0° to 90° can be seen as it appears from the bottom of the pump assembly.

**Movie S3. Close-up of microtissue lifting.**

Close-up of the capillary approaching several microtissues in sequence, showing how each microtissue is lifted off the surface and then transferred to a far location (off the screen).

**Movie S4. Demonstration of the operation of the robotic platform.**

(Bottom left) The capillary approaches a chosen tissue sample in a culture dish filled with microtissues. The tissue sample is picked up and transferred to the target 96-well plate (top left). A view from the side (right) is included.

**Data S1. (separate file)**

Rollerless eccentric peristaltic pump design file (.dwg).

**Data S2. (separate file)**

Rollerless eccentric peristaltic pump design and assembly descriptions (.pdf).
